# LBR nucleoplasmic domains regulate X-chromosome solubility and nuclear organization

**DOI:** 10.64898/2026.03.30.714681

**Authors:** Jonathan Fiorentino, Irene Perotti, Nerea Ruiz Blanes, Valentina Rosti, Ioanna Sigala, Eleni Nikolakaki, Alessio Colantoni, Annunziata D’Elia, Roberto Massari, Ferdinando Scavizzi, Marcello Raspa, Michela Ascolani, Neil E Humphreys, Thomas Giannakouros, Mitchell Guttman, Chiara Lanzuolo, Gian Gaetano Tartaglia, Andrea Cerase

## Abstract

The nuclear lamina plays a central role in genome organization, yet how specific lamina-associated proteins regulate chromosome architecture during development remains unclear. Here, we show that the nucleoplasmic domains of the Lamin B Receptor (LBR) are essential for X-chromosome localization at the nuclear periphery and chromatin architecture during neural differentiation. Using genetic dissection of LBR function, combined with genome-wide chromatin solubility profiling and transcriptional analyses, we demonstrate that loss of LBR N-terminal domains impairs proper cell differentiation and X chromosome inactivation (XCI), selectively disrupting chromatin structure in neural progenitors but not in pluripotent cells. Strikingly, these effects are disproportionately concentrated – but not limited to - on the inactive X chromosome, which undergoes a pronounced shift toward a more soluble chromatin state. Our findings establish the nucleoplasmic function of LBR as a key determinant of X-chromosome functionality and identify chromatin solubility and accessibility as a previously underappreciated layer of genome regulation by the nuclear lamina in XCI. Finally, our work provides definitive genetic evidence that LBR’s nuclear architectural functions are molecularly separable from its metabolic sterol reductase activity, which is preserved in our model, and are critically necessary for XCI in differentiating mouse female XX ESCs models.

## Introduction

X chromosome inactivation (XCI) is an intricate and evolutionarily conserved epigenetic process that ensures dosage compensation between male and female mammalian cells^1,2^. Initiated during early embryonic development, XCI results in the transcriptional silencing of one of the two X chromosomes in females, leading to the formation of a mostly transcriptionally inert structure known as the Barr body^3–6^. This fascinating mechanism allows for the equalization of X-linked gene expression between sexes, thereby preserving genomic balance.

While the molecular events regulating the onset^7–21^ and perpetuation of XCI have been extensively studied^22–26^, several studies have unveiled a new layer of complexity associated with the spatial organization of the inactive X chromosome (Xi) within the three-dimensional nuclear architecture^27–29^. The position of the Xi in the nucleus, particularly its tendency to reside at the nuclear periphery^27,28^ or in the proximity to the nucleolus^28^, has emerged as an important determinant in the establishment and/or maintenance of gene silencing^27,28^. Lamin B receptor (LBR) has been reported as a direct Xist RNA interactor and critical for Xist spreading into active genes and for its silencing function and the peripheral localization of the Xi^27^. This spatial arrangement is not merely a consequence but appears to actively contribute to both the initiation and the maintenance of transcriptional silencing^27–29^. Importantly, the Xi’s peripheral localization is not static but subject to alterations during different stages of the cell cycle and cellular differentiation^28,30^. The dynamic nature of nuclear positioning implies functional consequences. Indeed, the perinuclear environment offers a unique landscape enriched with heterochromatin-associated factors and repressive chromatin modifications, contributing to the consolidation of gene silencing on the X^27–29^, possibly through phase-separation mediated events^31–36^. Understanding the inticate dynamic spatial relationships provides crucial insights into the epigenetic regulation of X-linked genes and their impact on cellular function, both in health and disease. Nevertheless, while the molecular composition of the nuclear lamina has been extensively characterized, the specific mechanisms by which individual lamina-associated proteins regulate chromosome-specific, higher-order architecture during development remain incompletely understood^37–39^.

In this context, we investigated the role of LBR nucleoplasmic N-terminal domain, previously reported to interact with Xist RNA, in X chromosome inactivation and development. We employed recently-generated mouse and ES cells lacking the LBR N-terminal nuclear domains, a model that elegantly allowed us to separate the nucleoplasmic functions of LBR from its inner nuclear membrane (INM) associated sterol reductase activity. We show that, although the absence of the LBR N-terminal nucleoplasmic domain results in Pelger-Huet (PH) nuclear defects, such as chromatin clamping in white cells^27–29^, it does not cause any skeletal alterations in mice, indicating very distinct roles for the nucleoplasmic N- (RNA binding and heterochromatin organization roles) and the membrane embedded C-terminal domains (sterol reductase). Consistent with previous observations, we detect only very mild gene expression changes *in vivo*, with substantial sex-differences in the mouse model. By contrast, in differentiating female ES cells, we demonstrate that: (i) LBR is essential for proper X chromosome inactivation and gene silencing in differentiating female cells; (ii) LBR is required for correct neuronal gene expression in mouse neuronal cell progenitors (NPCs); and (iii) LBR N-terminal domains are necessary for proper nuclear chromatin organization and chromatin solubility of the Xi and to a lesser extent of other chromosomes.

## Results

### Lbr NT-KO adult mice exhibit sex-specific gene expression differences

The Lamin B Receptor (LBR) is an inner nuclear membrane protein with dual functionality: its nucleoplasmic N-terminal domain mediates interactions with chromatin, heterochromatin proteins and RNA, while its C-terminal domain encodes a sterol reductase enzyme essential for cholesterol biosynthesis^40–42^. LBR is ubiquitously expressed and plays critical roles in nuclear envelope integrity, chromatin organization and gene regulation during development^43,44^. In humans, LBR has emerged as an important disease gene, with mutations causing a spectrum of conditions ranging from the relatively benign Pelger-Huët (PH) anomaly — characterized by hypolobulated granulocyte nuclei — to the lethal Greenberg skeletal dysplasia (GB), depending on the nature and zygosity of the mutation^45–47^. Mutations in the LBR gene have also been associated with a wide spectrum of phenotypic abnormalities, including specific skeletal defects (other than GB) observed during fetal development such as syndactyly or polydactyly and/or alteration of long bones^48^ as well as perinatal dysplasia^49^. Notably, homozygous LBR N-terminal mutant mice missing its nucleoplasmic Tudor domain, the RS domain and a large part of the globular domain (NT-KO, Lbr236/Lbr236^29^) (**Fig. S1A**) show Pelger-Huet-like phenotypes^43,44,47,50,51^, but do not exhibit previously reported skeletal malformations, such as syndactyly or polydactyly^29^. To systematically assess potential skeletal phenotypes, we conducted an in-depth comparative analysis of the entire skeleton in homozygous LBR mutants and wild-type (WT) mice. Initial anatomical examination revealed no major skeletal difference between WT and LBR NT-KO mice. We decided, however, to evaluate potential minor alterations in rib cage and long bone development, particularly in adult mice, as previously reported from other groups^52^. To this end, we performed microtomography analysis of WT and LBR NT-KO mice. Our analysis revealed no statistically significant differences in the length of all bones examined and in the ribcage radius, supporting the hypothesis that homozygous Lbr236/Lbr236 mutants (LBR NT-KO) do not manifest any observable skeletal abnormalities. This suggests that skeletal defects are most-likely linked to C-terminal mutations and to compromised sterol reductase activity^41,44,46,53^ (**Fig. S1B-C**), clarifying a long-standing debate in the field of human genetics, linking Pelger-Huet and Greenberg dysplasia (GD) solely to the zygosity of the mutation (heterozygotic mutation associated to PH and GD to the homozygotic state, respectively)^45^.

In order to have a definite answer on the extent of gene deregulation *in vivo* in these animals, we extended the panel of RNA-seq samples from WT and NT-KO mice reported by Young et al.^29^ (see Material and Methods). Our analysis confirmed previous results (**Fig. S2 and Supplementary Table 1**). Specifically, we report higher gene expression differences in the male and only minimal deregulation in female samples, including limited changes in X-linked gene expression when comparing WT control and LBR NT-KO mouse models, in line with previous analyses^29^.

GO-term enrichment analysis in females did not return any statistically significant GO terms, while for upregulated genes in males the following GO terms were reported (**Supplementary Table S1**): oxidoreductase activity, monooxygenase activity, long-chain fatty acid metabolic process, likely linked to LBR reduction in the mutant animal, as previously reported^29^. No significantly downregulated GO terms were found in males.

### Xi localization and Barr body characteristics are altered in LBR NT-KO differentiating cells

Given the known role of LBR in Xi localization, we assessed whether the lack of interaction between Xist and LBR^27^ could be linked to changes in the Barr body characteristics. We used 2 independent female LBR NT-KO ES clones (A8 and B3) for the analysis, showing similar results. We first performed H3K27me3 immunofluorescence—used as a marker of the Barr body^54^ — upon NPC (neuronal progenitor cell) differentiation to confirm that LBR wild-type and LBR NT-KO cells harbours a comparable number of Xi prior to conducting downstream experiments **(Fig. S3)**.

We did not observe any differences in the percentage of cells having a Barr body between LBR NT-KO and control cells at 5 days of neuronal differentiation towards neuronal progenitor cells (NPCs), as determined by H3K27me3 immunofluorescence analysis, an Xi surrogate marker (**Figure S3A-B)**. Likewise, morphological examination of the Xi showed no notable or little differences in the circular shape of the Barr bodies. However, quantitative measurements of area, solidity, and mean fluorescence intensity further revealed a decrease in these parameters in mutant cells relative to WT cells, suggesting a potential alteration in the Barr body morphology and organization/function of the Xi (**Figure S3A-B)**.

### Peripheral localization of the Xi is compromised in LBR NT-KO NPCs

The identification of the lamin B receptor (LBR) as a binding partner of Xist^27^ suggests a pivotal role for LBR in anchoring X-linked regions to the nuclear periphery. This proposed function implies that the nuclear periphery environment facilitated by LBR may potentiate Xist RNA-mediated silencing, with LBR also being crucial for the initial spreading of Xist RNA onto active genes^1^. We therefore tested for the potential of our mutant to correctly localise to the inner nuclear membrane (INM) **(Fig. S4)**.

To assess whether the NT-KO–truncated LBR retains correct subcellular targeting, we performed a systematic localization analysis using GFP-tagged LBR expression constructs. Specifically, we assessed whether our mutant protein remained competent to traffic to and accumulate at the inner nuclear membrane (INM). To this end, we generated a panel of GFP-tagged LBR expression constructs in which distinct N-terminal and transmembrane regions were selectively preserved or removed, enabling us to dissect the contribution of individual domains to LBR targeting (**Fig. S4**). Among these constructs, the TMDLBR-GFP variant most closely recapitulates the architecture of our endogenous NT-KO allele. This mutant preserves all the multi-pass transmembrane region and sterol-reductase domain, while lacking the entire N-terminal chromatin-interacting region (**Fig. S4B**). Imaging analysis revealed that TMDLBR-GFP showed enhanced accumulation within the endoplasmic reticulum (ER), indicating partial retention along the secretory pathway. However, a substantial fraction of the protein clearly localized to the nuclear periphery, forming a clear and properly well-defined INM-associated rim (**Fig. S4B**) consistent with the localization pattern previously reported for comparable N-terminal LBR deletions^55^. These observations indicate that the N-terminal deletion reduces the efficiency of trafficking toward the INM but does not abolish targeting to the INM. Indeed, our truncated protein is still competent for membrane insertion and sterol reductase activity, although with increased ER residence time^40,56^. On the contrary, the mutant carrying the entire N-terminal domain and only the first transmembrane domain (LBRNtTMD1-GFP) localizes to the INM as previously reported^40^ (**Fig. S4B**. On the other hand, the LBR sterol reductase ortholog *TM7SF2*, completely fails to localize at the INM and accumulates exclusively at the ER, while the LBRNtTM7SF2-GFP (LBR N-terminal domain fused to TM7SF2) also localizes to the ER without a nuclear rim (**Fig. S4B)**. Taken together, comparison across the full construct panel further supports a cooperative requirement for both the N-terminal domains and the C-terminal transmembrane region: the latter is essential for membrane insertion and INM access, whereas the former contributes to efficient retention and stabilization at the nuclear envelope and nuclear lamina-ER shuttling^40^. Once we assessed the correct localization of our mutant protein to the INM, we decided to re-analyse in different ways, the frequency of Xi-lamina contact using microscopy in our mutant and matched controls cells in cell differentiation experiments, using different approaches.

We differentiated our cells to neuronal progenitors (NPCs) for 5 days (**Fig. S5A**) as previously described^29^. To evaluate the spatial distribution and characteristics of the inactive Xi foci in two cell lines, WT and LBR NT-KO, we conducted an analysis using concentric nuclear circles automatically drawn from the nuclear periphery to the center, defining up to three zones—zone 1 being the closest to the periphery and zone 3 the furthest (**Fig. S5B**) (see Materials and Methods for details). This analysis revealed a lower proportion of cells in the close proximity pf the nuclear lamina (zone1) in the LBR NT-KO cells compared to the WT cells (**Fig. S5C**). These results suggest that the Xi tends to be closer to the nuclear lamina in the WT cells than in the LBR NT-KO cells, in agreement with previous reports by Young et al and Chen et al^27,29^.

### Differentiating Lbr NT-KO female ES cells show significant XCI defects by bulk and scRNA-seq analysis

To test the extent of XCI defects at transcriptome-wide level in Lbr NT-KO female ESCs, we performed bulk RNA sequencing analysis on cells differentiated into neuronal progenitor cells (NPCs), using two independent clones (A8 and B3). First, we performed a Principal Component Analysis (PCA) and we computed sample-to-sample distances, which show that the samples are homogeneous for each condition (Lbr NT-KO and WT), while they cluster by condition both in mESCs and in NPCs at day 5 (**Supplementary Figure S6A-B**), suggesting the presence of extensive transcriptomic differences between the WT and the mutant cells.

We next performed a differential expression analysis between Lbr NT-KO and WT samples at both stages (see Materials and Methods). In Lbr NT-KO cells, we identified in the A8 clone 300 upregulated and 406 downregulated genes in mESCs, and 681 upregulated and 828 downregulated genes in NPCs (similar results were obtained in the B3 clone, **Supplementary Figure S7A** and **Supplementary Table S2**). Unexpectedly, in both cell types, the number of downregulated genes exceeded that of upregulated ones (**Supplementary Table S2)**. While LBR has been primarily associated with heterochromatin organization, this observation may be consistent with a scenario in which disruption of a structural component of nuclear architecture weakens domain boundaries, allowing repressive chromatin features to spread into transcriptionally active regions (see also Discussion). Importantly, a highly significant overlap was observed between mESCs and NPCs for both upregulated (87 genes, Fisher’s exact test p-value = 3.0 × 10^-61^) and downregulated genes (77 genes, Fisher’s exact test p-value = 4.6 × 10^-31^). Upregulated genes included transcriptional regulators and signalling components associated with early neural and cell fate specification, such as Otx2, Sp5, Bhlhe40, and Etv4, alongside mediators of Nodal and Wnt signalling (Lefty1, Lefty2, Wls) and cell adhesion (Cdh2, Grik3), while downregulated genes were enriched for factors linked to X-chromosome inactivation (Xist), pluripotency maintenance (Dppa3, H19, Peg10), and imprinted gene regulation (Klhl13, Mid1, Gadd45g, Shtn1, Rab6b, Ptk2b, Ephx2, Pdpn). These results are potentially indicating that a core set of Lbr-dependent transcriptional changes is conserved across differentiation stages.

We also investigated potential XCI defects in NPCs Lbr NT-KO vs control samples. In brief, we found that genes known to escape XCI^29^ are generally not differentially expressed, except for *Xist* and *Mid1 (*clone A8; Xist only clone B3), both of which are downregulated in Lbr NT-KO samples (**Figure 1A**). In contrast, differentially expressed genes (DEGs) are enriched for genes of chromosome X and, to a lesser extent, of chromosome 3 (**Supplementary Figure S6C**; see Materials and Methods).

**Figure 1.**
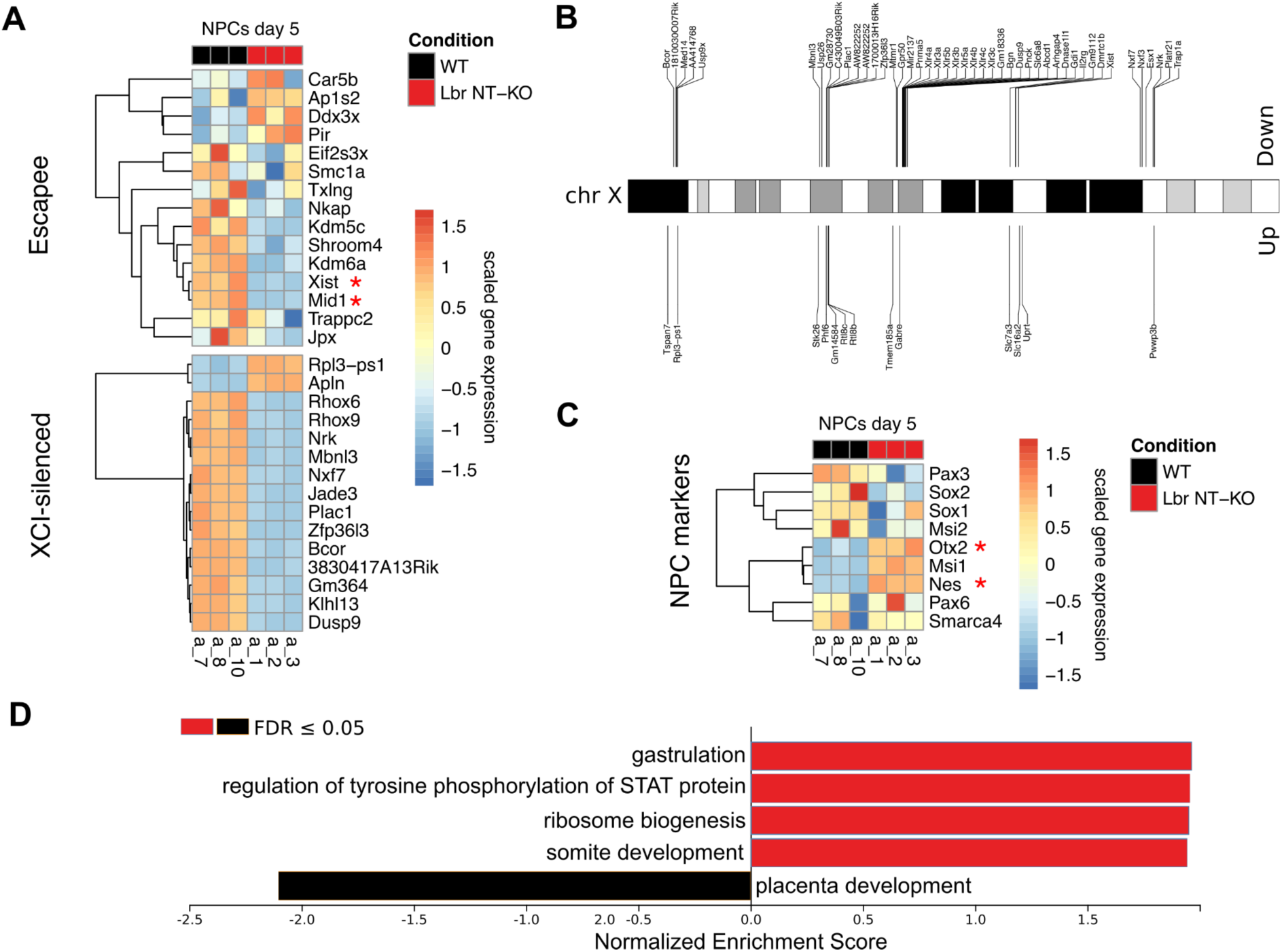
Bulk RNA-seq analysis of Lbr NT-KO and WT mESCs and NPCs at day 5 of differentiation. **A)** Heatmaps showing the expression of 16 genes known to escape X chromosome inactivation and the top 15 XCI-silenced genes differentially expressed between WT and Lbr NT-KO in NPCs at day 5. *Mid1* (log2FC= -3.05; FDR=10^-68^) and *Xist* (log2FC= -2.85; FDR=10^-44^) are differentially expressed escapee genes. **B)** Schematic representation of the X chromosome highlighting DEGs between WT and Lbr NT-KO. We binned the X chromosome (5Mb width) and we kept only bins with more than 5 DEGs. We show genes downregulated and upregulated in Lbr NT-KO samples at the top and bottom of the chromosome, respectively. **C)** Heatmap showing the expression of NPC marker genes in NPCs at day 5 of differentiation. *Otx2* and *Nes* are differentially expressed between WT and Lbr NT-KO samples, with log2FC= 1.74, FDR=10^-28^, and log2FC= 1.03, FDR= 10^-41^, respectively. **D)** Weighted set cover bar plot of the Gene Ontology (GO) terms related to Biological Process from a Gene Set Enrichment Analysis on the DEGs between WT and Lbr NT-KO in NPCs at day 5. We show the significant GO terms enriched in DEGs upregulated (red) or downregulated (black) in Lbr NT-KO samples. Analysis from the A8 data is shown.

In NPCs we found 119 (A8) and 104 (B3)/1509 (∼6-7%) DEGs belonging to the X chromosome, ∼ 1,5-2% more than what would be expected by chance given the size and gene content of the mouse X chromosome, with a preference for genes downregulated in Lbr NT-KO samples **Supplementary Figure S7B**). Notably, we found that these DEGs belong to specific regions (clusters) of the X chromosome supporting the idea that these regions might lose the correct conformation/structure (**Figure 1B**). Moreover, some of these regions are enriched for specific classes of repeat-associated and low-complexity sequence annotations (**Supplementary Figure S7C**), such as simple repeats (SR), low-complexity regions (LCR), long terminal repeats (LTR) and rolling circle (RC) repeats.

Together, these analyses suggest that LBR NT-KO in differentiating mESCs induces extensive transcriptomic changes, potentially impacting X chromosome inactivation initiation and maintenance^57^. Contrary to our expectations, we have more genes downregulated than genes upregulated upon loss of LBR nucleoplasmic domains. Differential expression analysis identified 82 downregulated and 37 upregulated genes, with the downregulated set notably enriched for X-chromosome inactivation-associated factors including *Xist*, members of the *Rhox* and *Xlr* gene families, and epigenetic regulators, while upregulated genes were predominantly associated with neuronal progenitor identity and neuronal signalling. Interestingly, most dysregulated genes belong to just 5 distinct regions of the X chromosome (**Figure 1B)**.

### Neuronal differentiation is affected in LBR NT-KO NPCs

Next, we looked at the expression of 9 known NPC marker genes in NPCs, and we found that *Otx2* and *Nes* are upregulated in Lbr NT-KO samples (**Figure 1C**), suggesting altered–potentially accelerated– differentiation dynamics in the mutant cells. We also performed a Gene Set Enrichment Analysis (GSEA)^58^ of the DEGs in NPCs at day 5 and we found that genes upregulated in Lbr NT-KO are enriched in Gene Ontology (GO) terms related to gastrulation and early development, whereas the downregulated DEGs are enriched in biological processes such as placenta development (**Figure 1D** and **Supplementary Table S3**). These results suggest that Lbr N-terminal domains may be needed for early development, which is in line with previous reports of elevated embryonic lethality in NT-KO LBR mice (**Fig. 1D and S12**)^29^. We also analysed if the differential progression can be attributed to specific cell subpopulations, by performing single-cell RNA-sequencing of Lbr NT-KO and WT NPCs at day 5 of differentiation (**Figure 2**). After quality control, we obtained 1410 and 2150 Lbr NT-KO and WT cells, respectively (**Supplementary Figure S8A**; see Materials and Methods).

**Figure 2.**
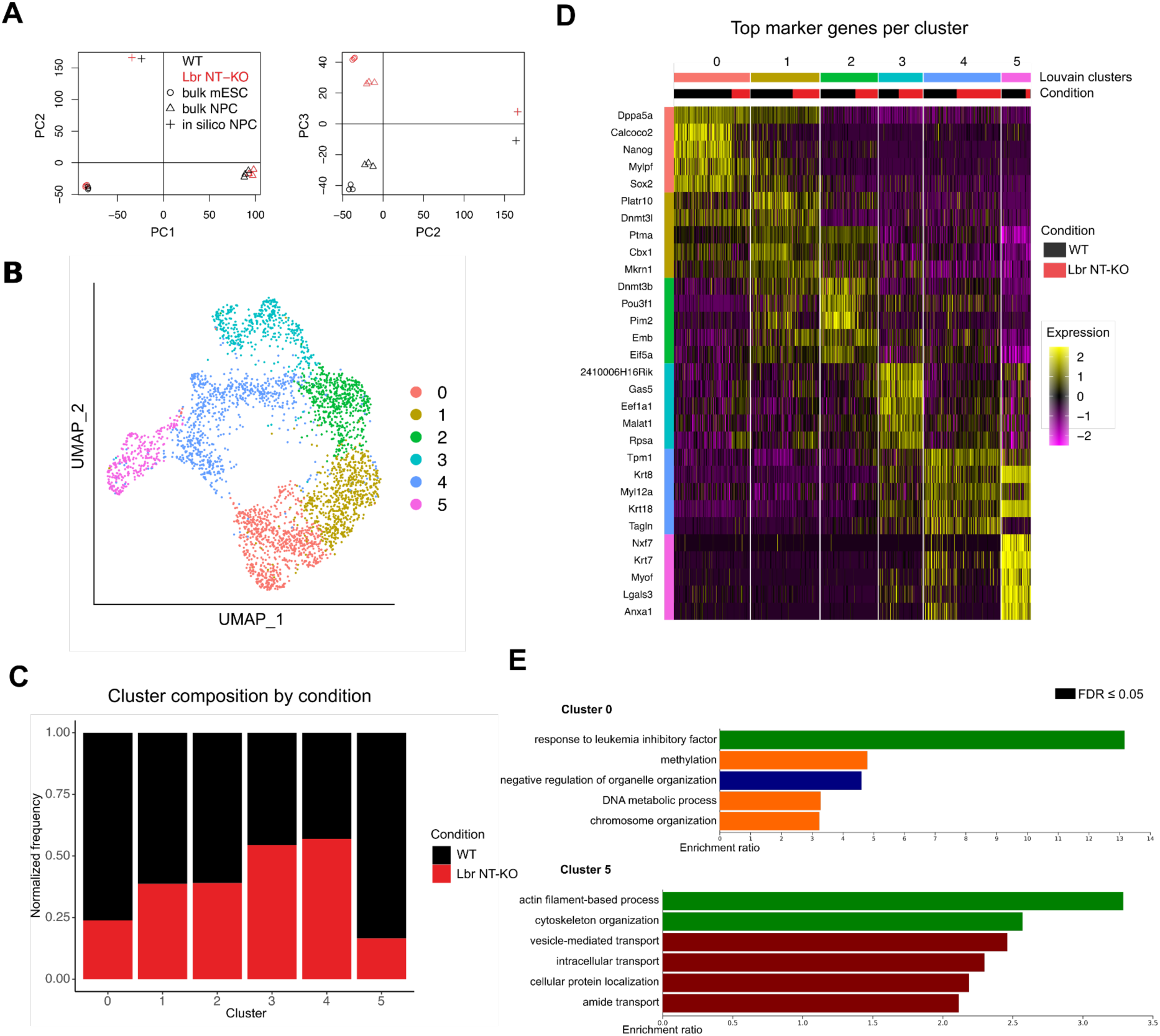
Single-cell RNA-seq analysis of WT and Lbr NT-KO neuronal progenitor cells (NPCs). **A)** PCA plot of integrated bulk (mESCs and NPC day 5, clone A8 shown) samples and scRNA-seq (NPCs day 5), indicated as “in silico NPC”, for WT and Lbr NT-KO **B)** UMAP plot showing cell clustering of WT and Lbr NT-KO NPCs. **C)** Bar plot of cluster composition by condition. On the y-axis the cell frequency normalized by the total number of cells by condition is shown. **D)** Heatmap displaying the top 5 marker genes per cluster, split by condition (see Methods for details on marker genes computation). **E)** Weighted set cover bar plots obtained from an Over Representation Analysis (ORA) of Gene Ontology Biological Process terms, for the set of marker genes of cluster 0 and cluster 5. Bar colors indicate functional categories: green, developmental processes; orange, chromatin and DNA-related processes; maroon, intracellular transport and localization; dark blue, organelle organization.

Integration and comparison of the bulk RNA-seq data with *in silico* bulk profiles computed from the single-cell data showed a strong correspondence of the differences between the mutant and WT samples in NPCs between data modalities or RNA-sequencing (**Figure 2A**; see Materials and Methods)^59^.

Next, we performed unsupervised clustering of the cells using the Louvain algorithm^60^ and we identified 6 cell clusters, which we visualized using a Uniform Manifold Approximation and Projection (UMAP) plot^61^ (**Figure 2B**). We found that most cell clusters were depleted for LBR NT-KO cells, while clusters 3 and 4 were slightly enriched for them **(Figure 2C**). We identified cluster-specific marker genes (**Figure 2D and Supplementary Table S4**) and we characterized each cell cluster by performing an Over Representation Analysis (ORA) on the specific marker genes (see Materials and Methods)^58^. We found a substantial heterogeneity between the biological processes characterizing the cell clusters, ranging from terms related to general metabolic processes and chromosome organization for clusters 0, 1, 2 and 3, to terms related to cell motility and cytoskeletal organization for clusters 4 and 5 (**Figure 2E, Supplementary Figure S8B and Supplementary Table S5).** Together, these results reveal the functional heterogeneity of NPC cell subtypes and that Lbr NT-KO cells are unevenly distributed across NPC subtypes, with enrichment in specific clusters and depletion in others. Given the subtlety of the transcriptomic differences between the cell clusters, and to quantify the presence of neuronal differentiation defects in Lbr NT-KO cells while taking into account the continuous nature of a cell differentiation process, we computed a diffusion map^62^ of all the cells and we inferred a differentiation trajectory using Slingshot^63^, finding two lineages (**Figure 3A**).

**Figure 3.**
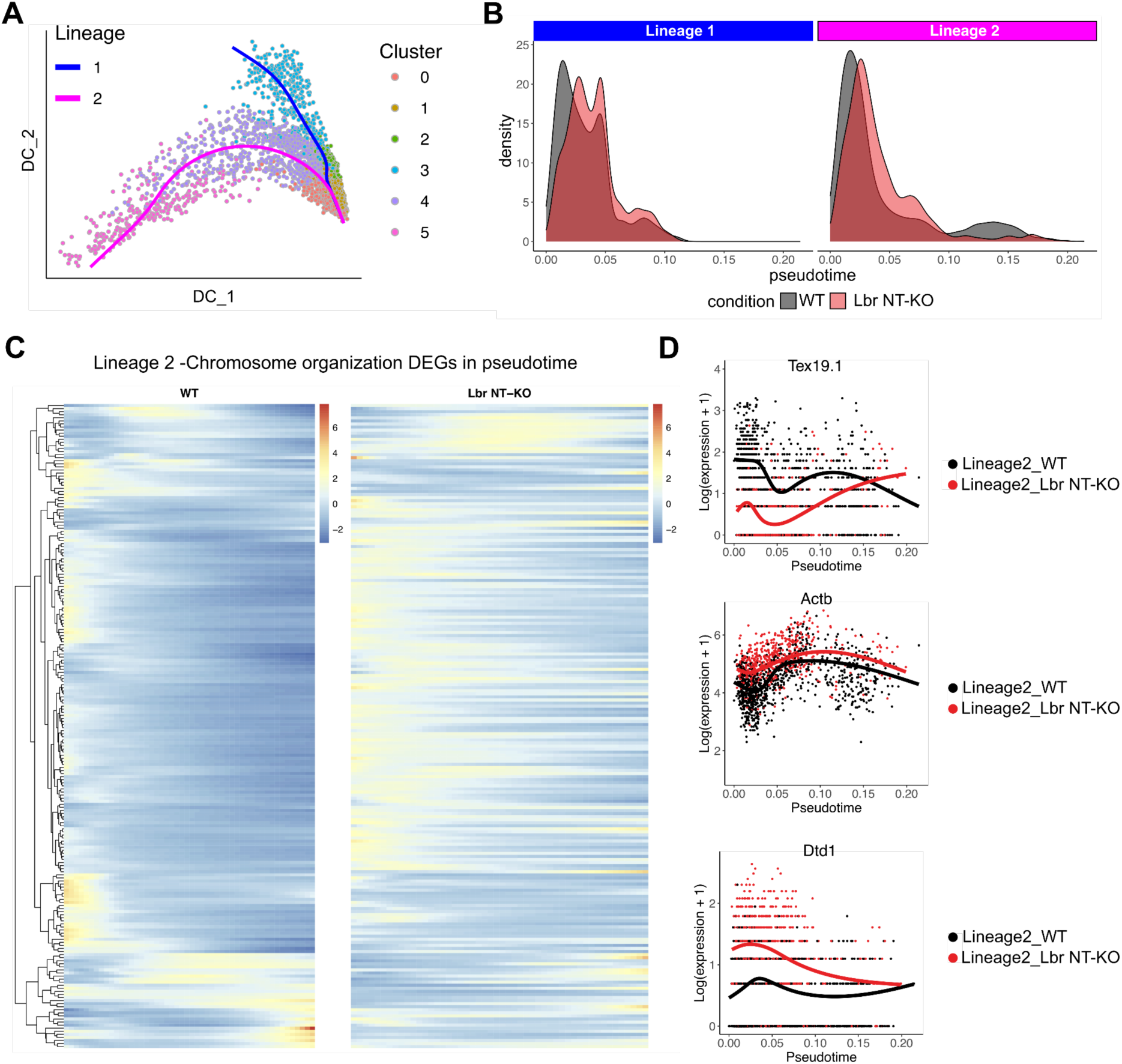
Trajectory inference and differential progression between Lbr NT-KO and WT cells. **A)** Diffusion map of WT and Lbr NT-KO cells colored by cluster. The two lineages inferred via Slingshot are overlaid to the diffusion map. **B)** Density plots showing the differential progression of WT and Lbr NT-KO cells along pseudotime for the two inferred lineages. **C)** Matched heatmap showing the expression of genes along pseudotime, in cells of lineage 2, belonging to the significant GO term “Chromosome organization” (see Figure S9 for the results of the functional enrichment analysis), from a functional enrichment analysis performed on genes with a differentially expression pattern between WT and Lbr NT-KO cells along lineage 2. **D)** Smoothers for the top three significant genes (according to the Wald test’s statistic) from panel C.

We chose cluster 0 as the root for the computation of the trajectory based on its higher expression of known naive pluripotency genes compared to the other clusters (**Supplementary Figure S9A**).

We independently confirmed our choice to assign cluster 0 as the least differentiated cluster by computing the initial macrostates of differentiation using CellRank^64^, an algorithm that integrates the RNA velocity^65,66^ with transcriptome similarity to study cell state dynamics (**Supplementary Figure S9B**; see Materials and Methods). We computed a pseudotime variable using Slingshot and we tested the overall differential progression along pseudotime between Lbr NT-KO and WT cells in the two lineages separately (**Figure 3B**; see Materials and Methods)^67^. We found a significant difference in both lineages (Lineage 1: p-value = 5.8 x 10^-11^; Lineage 2: p-value = 5.1 x 10^-10^), with Lbr NT-KO cells showing an overall advancement in pseudotime compared to the WT cells, except at the end of lineage 2, where Lbr NT-KO cells are depleted, in line with the observed cell cluster composition (**Figure 3B** and **Figure 2C**). To elucidate the differences at the gene level, we performed a differential expression test between the conditions along the two lineages using tradeSeq^68^. We found the most extensive differences between Lbr NT-KO and WT cells along lineage 2 (see Materials and Methods). Specifically, a functional enrichment analysis of the 3302 DEGs showed several significant GO terms, including “chromosome organization” (**Supplementary Figure S9C-D** and **Supplementary Table S6**; see Materials and Methods), with a range of patterns of differential expression between WT and Lbr NT-KO cells along the pseudotime (**Figure 3C-D**). These results suggest that Lbr NT-KO cells undergo an accelerated differentiation process, with a premature progression along both lineages compared to WT cells. However, their depletion at the end of lineage 2 indicates a potential failure to fully complete differentiation, which may reflect a loss of specific neuronal progenitor subtypes and/or X chromosome inactivation defects. Interestingly also polycomb repressive complex members such as EZH2, implicated in the regulation of proper cell differentiation and genome organization^69^, are deregulated between the Lbr NT-KO and wild-type cells. The differential expression analysis along pseudotime further highlights potential disruptions in chromosome organization, suggesting that epigenetic or transcriptional dysregulation may underlie the altered differentiation dynamics observed in Lbr NT-KO cells.

### Epigenetic changes predict gene deregulation in Lbr NT-KO cells

To examine the epigenetic landscape of genes dysregulated in Lbr NT-KO cells, we considered DEGs between Lbr NT-KO and WT samples from bulk RNA-seq data and we analyzed ChIP-seq data from neuronal progenitor cells^70^ (see Materials and Methods). Both downregulated and upregulated genes are localized in heterochromatin regions, depleted of active chromatin marks (H3K27ac, H3K36me3) and enriched of H3K9me3 and H3K27me3 marks, belonging to constitutive and facultative heterochromatin, respectively (**Supplementary Figure S10**). Interestingly, Ring1B binding discriminates downregulated and upregulated genomic regions, being enriched in regions upregulated and reduced in regions downregulated. This indicates that Lbr is involved in the regulation of a complex interplay between repressed, polycomb-regulated, chromatin domains.

To investigate at the genome-wide level how the loss of the N-terminal domain of LBR impacts higher-order genome organization, we employed 4f-SAMMY-seq chromatin solubility profiling^37,71,72^, which sequentially fractionates chromatin based on differential solubility and maps euchromatic and heterochromatic domains to their genomic coordinates by high-throughput sequencing, in WT and Lbr NT-KO, in undifferentiated ESCs and differentiated NPCs (see Materials and Methods). To validate that 4f-SAMMY-seq accurately reflects the chromatin accessibility landscape, we assessed whether the S2S/S3 solubility ratio — which represents the relative enrichment of soluble, accessible chromatin (S2S) over less soluble, compacted chromatin (S3) — correlates with established epigenomic marks of open and closed chromatin states. We could verify that genome-wide comparison of the S2S/S3 solubility ratio in WT NPCs with chromatin marks associated with open (H3K27ac, H3K4me3) and closed (H3K9me3) states^70^ indicated that 4f-SAMMY-seq faithfully captures accessibility features of eu- and heterochromatin (**Figure 4A**; **Supplementary Figure S11A**). As expected, the S2S-enriched, more soluble fraction is preferentially associated with active chromatin marked by H3K27ac and H3K4me3, whereas H3K9me3-marked, gene-poor heterochromatin shows lower S2S/S3 ratios, consistent with enrichment in the less soluble S3 fraction (**Figure 4A**).

**Figure 4.**
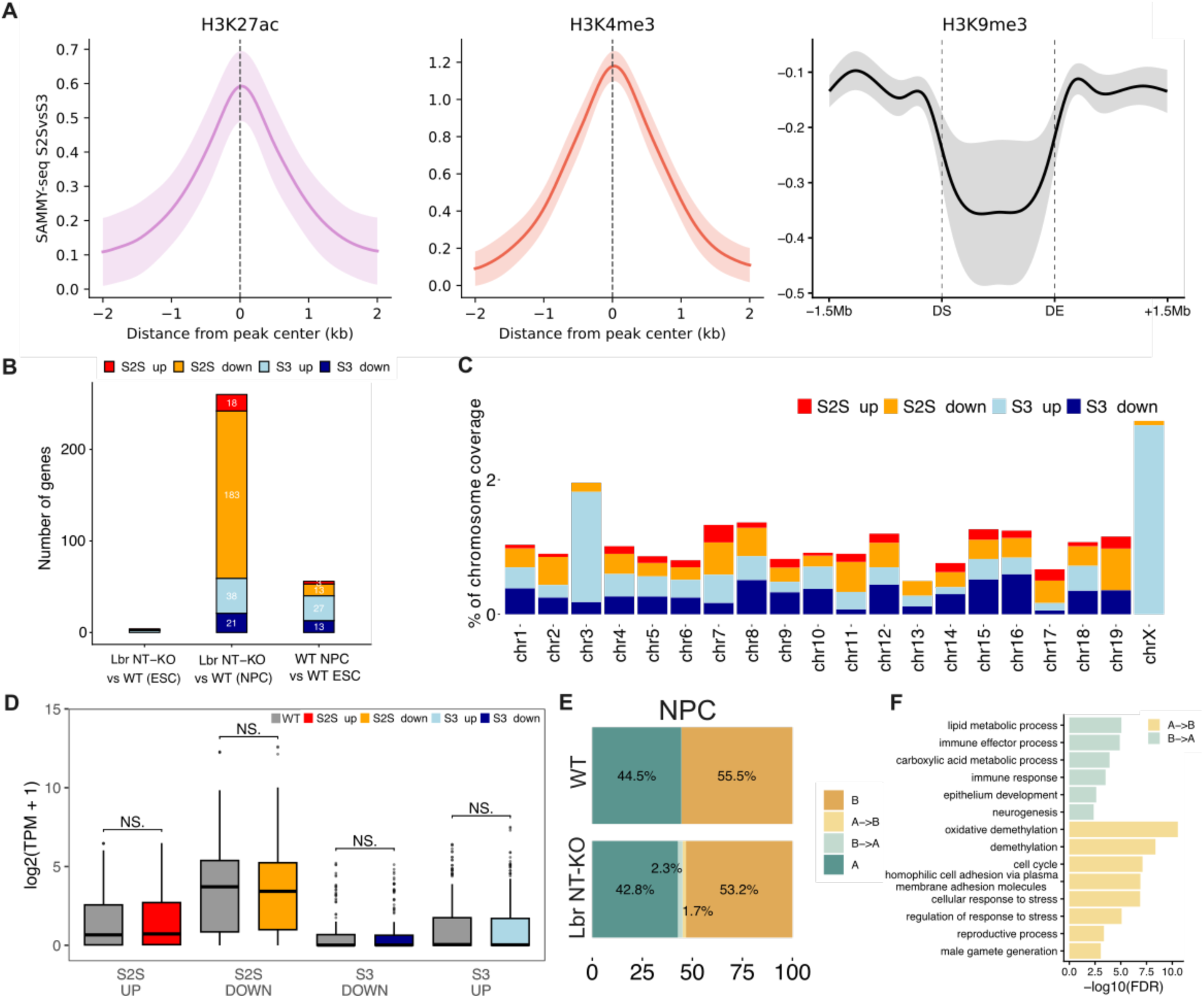
Integrated 4f-SAMMY-seq and RNA-seq analysis shows chromatin solubility defects and compartment switching in Lbr NT-KO NPCs. **A)** Metaprofiles showing the consensus WT SAMMY-seq enrichment ratio (S2S vs S3) in NPCs. Profiles for active chromatin marks (H3K27ac and H3K4me3) are centered on peaks. In contrast, the profile for the heterochromatin mark H3K9me3 is aligned to the boundaries of H3K9me3 domains, with the x-axis representing the relative position from the domain start (DS) to the domain end (DE), including 1.5 Mb flanking sequences on both sides. For each mark, the mean signal across biological replicates is shown, with shaded areas indicating the standard deviation. Positive values correspond to enrichment in the more soluble S2S fraction, whereas negative values indicate enrichment in the less soluble S3 fraction. **B)** Stacked bar plot showing the number of protein-coding genes with altered solubility (S2SvsS3). **C)** Box-plot distribution of log2 of transcripts per million (TPM + 1) in significantly differentially soluble regions in Lbr NT-KO (coloured) vs WT (grey) NPCs. The box lower and upper edges are the first and third quartiles, and the horizontal bar is the median. Whiskers extend up to 1.5 times the interquartile range (IQR) from the edges. Data points outside the range are outliers and are represented by dots. NS; Not significant (two-sided Wilcoxon test). **D)** Distribution of significantly differentially soluble regions across chromosomes in Lbr NT-KO in comparison to WT NPCs. **E)** Stacked bar plot of the average percentages of A and B chromatin compartments in WT, and compartments shifts in Lbr NT-KO. **F)** GO enrichment analysis of genes undergoing compartment changes in Lbr NT-KO NPCs. The bar plot represents significantly enriched and filtered biological process (BP) GO terms.

To capture statistically different soluble regions of eu- and heterochromatin we performed differential solubility analysis (see Materials and Methods). We found a stage-specific response to LBR N-terminal domains absence: ESCs displayed only minor solubility changes, whereas NPCs showed extensive remodeling, with a predominance of genes with reduced euchromatin accessibility (S2S down) (**Figure 4B).** GO analysis of S2S-down genes in NPCs revealed significant enrichment for vesicle-mediated and protein transport, intracellular signaling, and chromatin remodeling pathways, highlighting that reduced chromatin solubility preferentially affects genes governing trafficking, signal integration, and higher-order chromatin regulation (**Supplementary Figure S11B**). This is consistent with a model in which LBR-dependent anchoring contributes to the correct positioning and structure of chromatin in the nucleus; loss of LBR N-terminal interactions therefore causes chromatin alterations in euchromatin with a shift toward a more-soluble state. Across the NPC genome, altered solubility is distributed unevenly across chromosomes (**Figure 4C** and **Supplementary Figure S11E**). Notably, the X chromosome, and the chromosome 3 to a lesser extent, displayed an elevated enrichment of regions with increased S2/S3 fraction, indicating an accentuated shift toward a more soluble, less compact state. The specific remodeling of X chromosome structure confirms the altered observed transcriptional patterns (**Figure 1** and **Supplementary Figure S7C**) and strongly suggest a tailored action of LBR on X chromosome structure and function. We next examined transcriptional consequences of chromatin remodeling observed in the mutant cells using bulk RNA-seq, extracted in parallel to the 4f-SAMMY-seq. PCA of the bulk RNA-seq datasets confirmed clear genotype- and lineage-segregated transcriptional profiles, and the two bulk RNA-seq experiments show highly concordant log2(Fold Change) when comparing Lbr NT-KO vs WT (**Supplementary Figure S11C-D**). Genes associated with differentially soluble regions did not exhibit significant differences in TPM values between conditions, in both euchromatic and heterochromatic regions (**Figure 4D**). This indicates that alterations in chromatin solubility are not accompanied by consistent transcriptional changes. Interestingly, only one gene (Slc16a2) from the 5 X-linked DE clusters (**Fig. 1**) changed its solubility, indicating a general decoupling of gene-expression and chromatin solubility at this stage. This behavior is fully concordant with X-inactivation dynamics and highlights the sensitivity of the inactive X to perturbations in LBR-dependent nuclear architecture. From the comparison between the ESC (2Xa) and the NPC 1Xa, 1Xi we can parsimoniously assume that these differences likely and largely derive from the differential organization of the Xi in our mutant cells (**Supplementary Figure S11E),** but we cannot exclude limited changes from the Xa (see Discussion). To determine whether large-scale structural organization was also affected, we inferred A/B compartments in WT and mutant NPCs (see Materials and Methods). Compartment profiles indicated modest but clear rearrangements, with 2.3% of the genome shifting from B to A and 1.7% shifting from A to B in LBR NT-KO NPCs (**Figure 4E**). GO enrichment analysis uncovered a pronounced functional asymmetry in these transitions (**Figure 4F**). Genes relocating from A to B were strongly enriched for pathways linked to chromatin regulation, cell-cycle progression, oxidative demethylation, and stress responses, consistent with the collapse of regulatory programs upon loss of LBR-mediated nuclear organization. Conversely, B to A transitions activated metabolic pathways related to lipid and carboxylic-acid metabolism, alongside immune-related signatures and developmental processes such as neurogenesis and epithelial differentiation—potentially indicating that release from the nuclear lamina promotes metabolic rewiring and ectopic activation of immune and differentiation modules.

Together, these results demonstrate that the nucleoplasmic N-terminal domain of LBR plays a critical role in regulating chromatin accessibility, proper compartmental assignment, and transcriptional homeostasis. Its loss induces coordinated shifts in chromatin solubility, mainly affecting the inactive X chromosome, and compartment-specific transcriptional rewiring, particularly pronounced during neuronal differentiation.

## Discussion

The nuclear lamina is a key determinant of three-dimensional genome organization, yet the chromosome-specific roles of individual lamina-associated proteins during differentiation remain poorly understood. In this study, we dissect the nucleoplasmic function of LBR independently of its sterol-reductase activity and demonstrate that its N-terminal domains play a central role in stabilizing Xi architecture during neuronal differentiation^27,29^. Using a domain-specific genetic model, we show that loss of LBR nucleoplasmic domains does not impair skeletal development but leads to pronounced defects in nuclear positioning, higher-order chromatin organization, and chromatin solubility in differentiating female embryonic stem cells. These effects are most striking on the X chromosome, which displays a disproportionate shift toward a more soluble chromatin state in neuronal progenitor cells. Importantly, these architectural changes are only partially reflected at the transcriptional level. Our findings therefore position LBR as an architectural stabilizer of the inactive X chromosome, acting at the interface between nuclear positioning, chromatin solubility, and differentiation-dependent genome regulation.

Previous studies have linked LBR mutations to Pelger-Huet anomalies^43,44,47,50,51^ and to skeletal defects^41,44,46,53^, including polydactyly and syndactyly^29,43^ - primarily due to defects in cholesterol biosynthesis associated with its C-terminal sterol reductase activity -zygosity of the mutation. However, our data indicate that the N-terminal domains of LBR do not contribute to these skeletal defects, as LBR NT-KO mice exhibited no significant differences in bone length, rib cage circumference, or overall skeletal structure when compared to WT controls. This suggests that skeletal phenotypes observed in other LBR mutations may be primarily driven by disruptions in cholesterol metabolism rather than nuclear functions mediated by the N-terminal domain, and not simply linked to the zygosity state of the mutation. This work can also finally reconcile the differences between the mouse model of LBR mutation and the human pathology.

The role of LBR in XCI has been previously suggested, particularly in relation to its interaction with Xist and the organization of the inactive X chromosome (Xi) at the nuclear periphery. Our study confirms and extends these observations^27,29^, demonstrating that LBR NT-KO differentiating ES cells fail to properly localize the Xi to the nuclear periphery, leading to defects in gene silencing, chromatin organization and solubility. Despite this, LBR NT-KO adult female mice do not exhibit overt XCI defects, possibly due to compensatory mechanisms that emerge during development. The observation that only ∼30% of LBR NT-KO mice that make into adulthood (irrespectively of the sex) suggests that selection pressures may favor individuals in which alternative pathways ensure Xi localization (i.e. at the nucleolus) and silencing^28,29^. Indeed, in order to understand whether embryonic lethality occurred at early embryonic development stages and whether only female embryos were affected, we decided to perform immuno histochemistry analysis on e6.5 embryos (decidua stage) using H3K27me3 as a surrogate marker of the Xi (and embryo sex proxy). This analysis suggested that both male and female embryos get reabsorbed at decidua stage (independently of sex), indicating that Lbr has developmental roles either than XCI in females. This observation potentially indicates that a stochastic percentage of females that survive birth has achieved peripheral Xi localization in an LBR-independent fashion.

Our findings also reveal that LBR plays a critical role in neuronal differentiation, as LBR NT-KO neural progenitor cells (NPCs) display misregulation of key neuronal genes (**Fig 1 and 2**). This suggests that LBR contributes to chromatin organization necessary for proper neuronal gene expression. Our integrated analysis of chromatin solubility and transcriptional profiles further elucidates the impact of LBR N-terminal loss on higher-order genome organization. 4f-SAMMY-seq profiling revealed that Lbr NT-KO NPCs exhibit remodeling of chromatin solubility, with a pronounced reduction in the S2S fraction at genes involved in vesicle transport, signaling, and chromatin regulation, consistent with a relative shift toward less accessible chromatin fractions (**Fig. 5**). On the contrary, an increase in solubility was evident on the X chromosome, consistent with defective X-inactivation, and was accompanied by compartment-specific transcriptional changes, including A-to-B shifts at regulatory genes and B-to-A transitions at metabolic and developmental loci; and to a lower extent to chromosome 3, the two chromosomes carrying most DE genes (**Fig. S6C**). Interestingly, most gene silencing on the X is likely coming from the active X chromosome (Xa), possibly indicating that either LBR has a role in chromatin boundaries on all chromosomes, including the Xa or that reactivation of genes on the Xi(at sub-detection level) can trigger compensatory silencing events on the Xa. These findings indicate that the LBR N-terminal domain is essential not only for maintaining chromatin accessibility but also for proper compartmentalization, transcriptional homeostasis and XCI during neuronal differentiation, highlighting different roles for the lamina protein in the process of nuclear organization^37,71–73^.

**Fig. 5.**
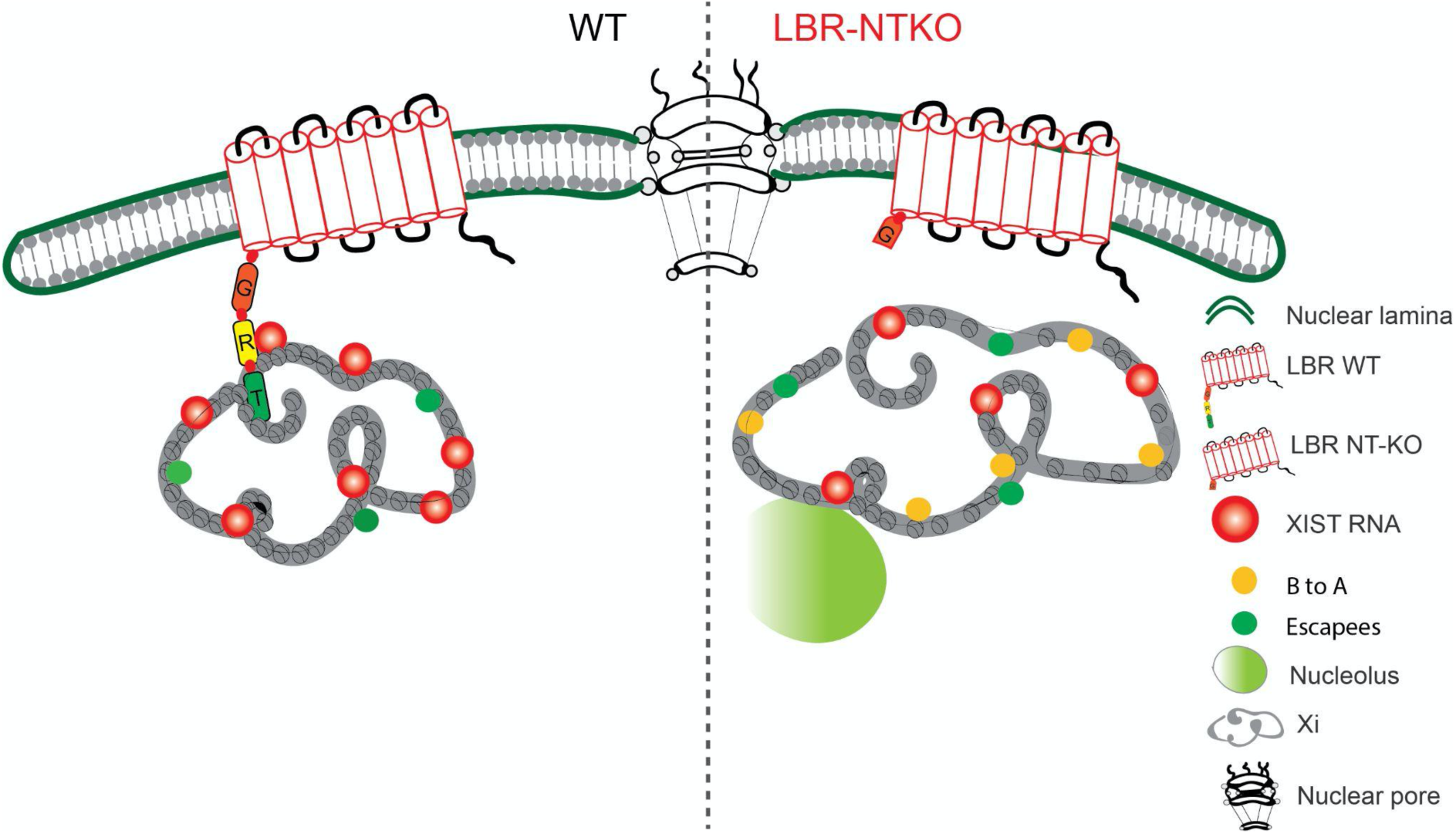
Changes in Xi-chromosome solubility are shown in the LBR-NT mutant NPCs. Left, In wild-type NPCs, functional LBR anchors the inactive X chromosome (Xi) to the nuclear periphery. The Xi is organized into two megadomains: the B compartment (decorated with Xist RNA) and the A compartment (containing escapee genes with RNA Pol II). Chromatin maintains normal solubility with a balanced S2S/S3 ratio. Right, Loss of LBR N-terminal domain disrupts Xi peripheral anchoring, causing the chromosome to relocate away from the nuclear membrane (possibly toward the nucleolus). The Xi becomes less compact with reduced inter-megadomain spacing and a pronounced shift toward a more soluble chromatin state (increased S2S/S3 fraction). Compatible with a decrease in compaction, more genes previously silent are now expressed (B to A transitions, yellow circles). Abbreviations: LBR, Lamin B Receptor; S2S, soluble chromatin fraction; S3, insoluble chromatin fraction; RNA Pol II, RNA Polymerase II. G, R and T represent, respectively, the Globular, the RS and the Tudor domain of LBR.

Differences between our results and previous studies examining LBR function in X-chromosome inactivation are likely attributable to the nature of the genetic perturbations employed^74^. In our study, LBR depletion selectively disrupts the nucleoplasmic form of the protein while leaving the sterol reductase domain intact, allowing us to uncouple LBR’s chromatin-organizing function from its essential metabolic role. By contrast, several prior studies^75^ relied on full LBR knockout models or shRNA/siRNA-mediated depletion strategies that eliminate or severely compromise the sterol reductase domain. Cholesterol biosynthesis is essential for membrane integrity and nuclear envelope function, and loss of LBR enzymatic activity cannot be compensated by TM7SF2 alone^76^, particularly during differentiation. As a consequence, complete LBR ablation is expected to introduce secondary defects in membrane composition, nuclear envelope stability, and cellular physiology, including chromatin organization confounding interpretation of chromatin and X-chromosome phenotypes^77,78^. Our domain-specific approach therefore reveals a direct and previously obscured role of the LBR nucleoplasmic domain in regulating X-chromosome architecture, independent of sterol metabolism. Importantly, earlier studies predominantly employed inducible *Xist* systems^75^, which do not recapitulate the physiological *Xist* downregulation seen in differentiating XX female cells, thereby masking defects in primary silencing and XCI organization.

Together, our results establish the nucleoplasmic N-terminal domains of LBR as essential regulators of inactive X chromosome nuclear positioning, chromatin solubility, and architectural stability during neuronal differentiation. By uncoupling LBR’s nuclear architectural role from its metabolic sterol-reductase function, we demonstrate that lamina-mediated regulation of chromatin solubility constitutes a distinct layer of genome organization that is particularly critical for the inactive X chromosome. The pronounced sensitivity of the Xi to loss of LBR nucleoplasmic domains, despite relatively modest transcriptional changes, reveals a buffering between chromatin material state and steady-state gene expression and highlights chromatin solubility as an underappreciated dimension of X-chromosome regulation. More broadly, our findings suggest that nuclear lamina components contribute to developmental gene regulation not solely by enforcing transcriptional repression, but by stabilizing higher-order chromatin architectures that ensure robustness of cell fate programs. Elucidating how LBR and other lamina-associated proteins cooperate with chromatin modifiers and phase-separated nuclear compartments will be an important avenue for future studies aimed at understanding genome regulation in development and disease.

## Author contributions

Andrea Cerase, Michela Ascolani, Neil Humphreys, generated and characterized the LBR-NT KO mouse and matched ES lines. Nerea Blanes characterized the A8 clone and conducted both the bulk and single cell RNA-seq experiments for clone A8. Jonathan Fiorentino performed all the bioinformatic analysis in this paper with the support of Alessio Colantoni. Annunziata D’Elia, Roberto Massari, Ferdinando Scavizzi, Marcello Raspa performed the microCT analysis of WT and LBR NT-mice. Irene Perotti performed the analysis of the microCT data, the characterization of clone B3. Irene Perotti and Valentina Rosti performed the differentiation experiments for the second bulk RNA datasets (clone B3) and for the 4f-SAMMY-seq analysis. Valentina Rosti performed the 4f-SAMMY-seq experiment with the support of Irene Perotti. Ioanna Sigala and Eleni Nikolakaki performed all the LBR localization experiments. Thomas Giannakouros, Mitchell Guttman, Chiara Lanzuolo and Gian Tartaglia provided expert advice and guidance and/or logistic or experimental support. Andrea Cerase, Jonathan Fiorentino and Irene Perotti wrote the paper and made all the figures. Andrea Cerase planned and conceived all the experiments performed in this paper.

## Acknowledgements

We would like to acknowledge: Giuseppe Trigiante for initial contribution to this work; Sara Bonomo, Gabriele Colozza, Giacomo Cavalli and Hegias Mira-Montebal for critical reading of this manuscript; Ugo Iannacchero and Emanuele Di Patrizio Soldateschi for advice in the analysis of 4f-SAMMY-seq data; all the Cerase, Tartaglia and Lanzuolo labs for their input, feedback and discussion; the QMUL Genome Centre facility for the 10X single cell analysis; Andreas Hierholzer, for helping deriving and characterize the early ES clones; all the EMBL Rome facilities where this work started, especially the mouse house facility; Emerald Perals and Monica Di Giacomo for embryo sections and staining and acquisitions. Vladimir Benes and all the GeneCore team; Andreas Buness for initial bioinformatics support; Luke Gammon for helping with the analysis of LBR-nuclear lamina distances, with the support of Nerea Blanes; Manish Kumar for support with RNA extraction from animal tissues; Kim Remnas from the Protein Expression and Purification Core Facility for kindly donating a tube of LIF that allowed some key experiments to be repeated exactly in the same experimental conditions. Finally, we would like to thank Phil Avner: without his help and support, this work would not have been possible.

## Declarations

Ethics approval and consent to participate. Ethical approval was not applicable. All authors have read and approved the manuscript before submission.

## Materials and Methods

### MicroCT acquisition and image morphometric analysis

High-resolution micro-computed tomography (microCT) was performed using a MILabs Hybrid OI/CT system (MILabs, Houten, The Netherlands). Acquisition comprised 720 projections over a 360° rotation (35 kV, 0.43 mA, 200 ms exposure per projection). Each scan lasted approximately 5 minutes and generated isotropic volumetric datasets for morphometric evaluation.

Bone length and rib cage geometry were quantified from reconstructed volumes. Linear bone measurements were obtained using the 3D measurement tool in Imalytics Preclinical 3.0 (Gremse-IT GmbH, Aachen, Germany). Rib cage radius was determined in ImageJ (version 1.54f, National Institutes of Health, USA) following segmentation of the body volume and calculation of the rib cage perimeter in a defined cross-sectional plane.

### Culture of mESCs

Mouse embryonic stem cells (mESCs) were cultured in 2i medium on 0.1% gelatin-coated flasks and passaged using Accutase upon reaching approximately 80% confluency. The experiments were performed using LBR NT-KO ES cell clones (A8 and B3), which were previously generated as described in PMID: 33846535.

### Differentiation of mESCs towards NPCs

For neural induction in vitA-deficient medium, 300,000 mESCs were plated onto laminin-coated 6-well plates in neuronal differentiation media composed of DMEM/F-12 (Life, 31331-028) and Neurobasal-A Medium (Life, 10888-022) 1:1, supplemented with B-27 Supplement minus vit. A (Life, 12587-010), N-2 Suppl (Life, 17502-048), BSA 25mg/mL (Sigma, A3311-10G), GlutaMAX Suppl. (35050-038), Insulin 20mg/mL (Sigma, I1882-100MG), 1-Thioglycerol (Sigma, M6145-25ML), Heparin 50 mg/mL (Sigma, H3149-10KU), Pen-strep (Life, 15140-122), 1% Fetal Bovine Serum, and 1:20000 FGF-basic (Preprotech, 450-33). Media was changed daily with fresh FGF-basic for a total of 5-6 days.

### Single-cell RNA sequencing preparation

Cell monolayer was washed twice in PBS, incubated in accutase for 5-7 minutes at 37 °C and neutralised with 10% fetal bovine serum and 0.2% chicken serum diluted in PBS. Harvested cells were centrifuged at 700 g for 5 minutes and cell pellets were washed with PBS, centrifuged, and washed again in PBS before passing the cells through a 0.10 µM filter. The cell suspension was then centrifuged at 700 g for 5 minutes and the pellet resuspended in 1 mL of 2% BSA diluted in PBS.

### Immunofluorescence

Cell monolayer was washed in PBS, fixed in ice-cold 4% PFA for 10 minutes, permeabilised with 0.5% Triton-X for 5 minutes and incubated for 1 hour at room temperature in blocking buffer composed of 1% BSA diluted in PBS. Incubation with primary (H3K27me3, 1:750; LBR, Active Motif #39536, 1:500 or Lamin B1, 1:500, Proteintech #66095-1-Ig) and secondary antibodies diluted in blocking buffer was kept for 1 hour at room temperature, followed by three 2-minute washes in PBS. Coverslips were mounted on DAPI-containing mounting medium.

### Image analysis

Image cell analysis was performed in ImageJ (Fiji 2.14.0/1.5f). The images of WT cells were taken with a magnification of 63X while LBR NT-KO were taken with a magnification of 40X. Thresholding was applied to images highlighting the signals to be measured by choosing “Max Entropy”. Area, solidity, and circularity were calculated from those thresholded areas using the specific tool “analyze particles” in ImageJ setting the dimension of the particles between 9 and infinite and the circularity between 0.02 and 1.00 to avoid the measurement of filamentous structure. To normalize the values of the Area of the WT Barr bodies from the particle analysis all the values obtained from the WT images were multiplied by the square of the normalization factor given by the ratio between 40 and 63 which represent the two magnifications at which the images were acquired. To remove outliers, all values above and below twice the mean were removed.

### Immunofluorescence and Analysis (Barr bodies)

To identify individual cell nuclei, cells were stained with DAPI and visualized under a fluorescence microscope. The inactive X chromosome (Xi) was detected using immunofluorescence staining for H3K27me3, while the nuclear periphery was marked by staining for Lamin B1 protein. For spatial analysis, all images were processed using ImageJ. Initially, each image was converted to RGB format. A straight line was manually drawn from the center of the Xi to the nearest point on the Lamin B1-stained nuclear periphery. The “RGB Line Profile” plugin in ImageJ was employed to generate an intensity profile along the drawn line, and distances were measured based on the peak intensities in the green (Lamin B1) and red (H3K27me3) channels. In cases where the fluorescence signal from Lamin B1 overlapped with the H3K27me3 signal, the distance was recorded as zero.

### Localization analysis

#### LBR/TM7SF2 constructs

Total RNA (1 μg), that was isolated from human testis using the TRIzol reagent, was converted to cDNA using M-MLV Reverse Transcriptase (Invitrogen) and random hexamer primers, according to the manufacturer’s recommendations.

*LBR-GFP:* Full-length LBR was amplified with PCR from 5% of the first strand reaction (1 μl) with the upstream primer LBRsense 5’-GATCTCGAGATGCCAAGTAGGAAATTTGCCGAT-3’ (carrying an additional XhoI site at its 5′ end) and downstream primer LBRantisense 5’-TGGATCCCGGTAGATGTATGGAAATATACGGTAG-3’ (carrying an additional BamHI site at its 3′ end) and ligated into the pGEM-T Easy vector (Promega). The insert was then removed from pGEM-T Easy, following digestion with XhoI/BamHI and ligated into the XhoI/BamHI site of pEGFP-N1.

*TMDLBR-GFP:* The cDNA expressing all the transmembrane domains of LBR, was amplified with PCR using full-length LBR as template with the upstream primer TMDLBRsense 5’-ATCTCGAGGAGTTTGGAGGAGTACCTGG-3’ (carrying an XhoI site at its 5′ end) and downstream primer LBRantisense and ligated into the pGEM-T Easy. The insert was then removed from pGEM-T Easy, following digestion with XhoI/BamHI and ligated into the XhoI/BamHI site of pEGFP-N1.

*LBRNtTMD1-GFP:* The plasmid expressing the LBR N-terminal domain and the first transmembrane domain fused with GFP was purchased from EMBL, Heidelberg (P30453, Ellenberg plasmids).

*TM7SF2-GFP:* TM7SF2 was amplified with PCR from total human testis cDNA with the upstream primer TM7SF2sense 5’-GATCTCGAGATGGCCCCCACTCAGG-3’ (carrying an XhoI site at its 5′ end) and downstream primer TM7SF2antisense 5’-TGGATCCCGGTAGATGTAGGGCATGATGCG-3’ (carrying a BamHI site at its 3′ end) and ligated into the pGEM-T Easy. The insert was then removed from pGEM-T Easy, following digestion with XhoI/BamHI and ligated into the XhoI/BamHI site of pEGFP-N1.

*LBRNtTM7SF2-GFP:* The following procedure was followed to make a construct expressing a hybrid protein in which the transmembrane domains of LBR were replaced with TM7SF2. TM7SF2 was amplified with PCR from total human testis cDNA as before but with primers without carrying the additional restriction sites (TM7SF2s 5’-ATGGCCCCCACTCAGGGCC-3’; TM7SF2as 5’- TCAGTAGATGTAGGGCATGATGCGGTAAG-3’). Two PCR reactions were subsequently performed; the first one with TM7SF2 as template, an upstream primer carrying at its 5’ end, a short sequence from the 3’ region of the N-terminal domain of the LBR cDNA (LBRTM7SF2commonsense 5’-AGTGAGAACCTTTGAAATGGCCCCCACTCAGG-3’) and

TM7SF2antisense; the second one with LBR as template, the upstream primer LBRsense (see above) and the downstream primer LBRTM7SF2commonantisense 5’-CCTGAGTGGGGGCCATTTCAAAGGTTCTCACT-3’. For the second reaction the template and the primers were denaturated at 95 °C for 5 min and then left overnight to hybridize. An initial extension was then performed at 68 °C for 30 min followed by 30 cycles with 1 min extension/annealing at 68 °C. The two PCR products were purified from the respective agarose gels, denaturated at 95 °C for 5 min and then left overnight to hybridize. A final PCR which included an initial extension at 68 °C for 30 min was then performed with the upstream primer LBRsense and downstream primer TM7SF2antisense. The product, LBRNtTM7SF2, was first ligated into the pGEM-T Easy vector and then removed, following digestion with XhoI/BamHI, and ligated into the XhoI/BamHI site of pEGFP-N1.

#### Transfection and Immunofluorescence Detection of GFP-tagged LBR constructs

HeLa cells were cultured in DMEM medium supplemented with 10% (v/v) fetal bovine serum (FBS) and antibiotics. Cells were incubated at 37 °C with 5% CO_2_. Transfections of LBR-GFP constructs were done with the Xfect™ transfection kit (Clontech-Takara Bio, Mountain View, CA, USA) according to the manufacturer’s instructions. Briefly, 4 × 10^4^ HeLa cells were plated on glass coverslips in 24-well plates, and 1 μg of plasmid DNA was diluted with Xfect Reaction Buffer to a final volume of 25 μl and added to 0.25 ml DMEM (without FBS). Following 4 h of incubation, nanoparticle complexes were removed via aspiration, and 2 ml of fresh complete growth medium was added. After 48 h, the cell coverslips were fixed with 4% paraformaldehyde in phosphate-buffered saline (PBS) for 20 min at room temperature, and excess aldehyde was quenched with 100 mM Tris-HCl pH 7.5. To visualize endogenous LBR, control cells were permeabilized with 0.2% Triton X-100 in PBS for 10 min, blocked with 0.5% fish skin gelatin (FSG) in PBS for 30 min and then probed with a primary anti-LBR rabbit polyclonal diluted 1:1000 (Novus, NBP2-56993) and secondary antibody (Alexa 488 goat anti-rabbit diluted 1:400). DNA staining of both control and transfected cells was performed with 1 mg/ml propidium iodide (PI) following treatment with 0.2 u/μl RΝase for 30 min. After 3x washing, the coverslips were mounted in 90% glycerol and visualized in a Nikon confocal microscope using the EZ-C1 3.20 software.

### Bulk RNA-seq analysis

Bulk RNA-seq data from 6 female (3 WT and Lbr NT-KO) and 6 male (3 WT and Lbr NT-KO) liver samples were processed as in Young et al^29^. Eight of these samples were already public at GEO (accession number GSE165447) while we added four samples. We performed a differential expression analysis, separately for female and male samples, between Lbr NT-KO and WT samples, using DESeq2 (version 1.30.1), controlling for the effect of the experiment in the design variable. Then we shrank the log2 fold changes using the function lfcShrink, with the apeglm estimator^79^. We defined differentially expressed genes (DEGs) as those having False Discovery Rate (FDR) < 0.05 and log2 fold change > 1 (in absolute value), as in our previous study^29^. In female samples, we obtained 4 DEGs upregulated in Lbr NT-KO samples and 4 downregulated; in male samples, we obtained 56 DEGs upregulated in Lbr NT-KO samples and 34 downregulated, in line with our previous results^29^.

For the bulk RNA-seq data from mESCs and NPCs at day 5, from clone A8, we have 12 samples in total, 6 Lbr NT-KO and 6 WT samples. RNA-seq reads were trimmed using the cutadapt software (version 4.1)^80^ to remove adapter sequences, setting the minimum read length after trimming to 18. We verified that the trimming was successful by inspecting the adapter content in the report obtained by running the FASTQC software (http://www.bioinformatics.babraham.ac.uk/projects/fastqc) on the trimmed fastq files.

Next, we used Bowtie2 (version 2.2.5)^81^ to align reads to rRNA sequences, since they were overrepresented in the report from the FASTQC software, and we removed reads mapping to these sequences. The remaining reads were aligned to the mouse genome and transcriptome (Ensembl GRCm38.98)^82^ using STAR (version 2.7.10a)^83^.

We performed a differential expression analysis between Lbr NT-KO and WT samples in mESCs and NPCs at day 5 using DESeq2 (version 1.30.1)^84^ and we shrank the log2 fold changes using the function lfcShrink, with the apeglm estimator^79^. We defined differentially expressed genes (DEGs) as those having False Discovery Rate (FDR) < 0.01 and log2 fold change > 1 (in absolute value). In mESCs, we obtained 300 DEGs upregulated in Lbr NT-KO samples and 406 downregulated; in NPCs, we obtained 681 DEGs upregulated in Lbr NT-KO samples and 828 downregulated. The lists of DEGs in mESCs and NPCs are provided in **Supplementary Table S2**. Next, we performed a GO term biological process Gene Set Enrichment Analysis (GSEA) using Web Gestalt 2019 through the R package WebGestaltR (version 0.4.5)^58^. We selected enriched categories using a p-value cutoff of 0.05. The enriched terms are reported in **Supplementary Table S3**.

For the analysis of chromosomal enrichment of DEGs, we computed the fraction of DEGs over the total number of genes tested for differential expression in each chromosome. Next, we built a contingency table for each chromosome and we tested the enrichment using a Fisher’s exact test, performed in Python using the function fisher_exact from the scipy package (version 1.7.2). Finally, we corrected the p-values for multiple testing using the Python function multiple_tests from the statsmodels package (version 0.13.1), with Benjamini-Hochberg FDR correction method (fdr_bh).

The Principal Component Analysis (PCA) plots were obtained using the function plotPCA from the DESeq2 package. Heatmaps and sample to sample distance matrices were obtained using the R package pheatmap (version 1.0.12). For the heatmaps displaying gene expression, we centered and scaled the expression level of each gene. The karyoplot was drawn using the R package karyoploteR (version 1.20.3)^85^. Next, we binned the X chromosome in regions of 5 megabases, and we retained five regions that contain at least 5 DEGs. We obtained the annotation for repetitive elements for GRCm38/mm10 from the UCSC Genome Browser^86^ and we counted the appearances of each type of repetitive element in each region. To compute an empirical p-value, we randomly sampled N=1000 non overlapping regions of the same length as the regions of interest and we repeated the count of the repetitive elements. In this way we built a distribution from which we computed the p-value as the probability of getting a value larger than that obtained from the region of interest by chance.

### Single-cell RNA-seq analysis

Alignment and counting were performed using the Cell Ranger pipeline (version 6.1.2, GRCm38 reference)^87^ for the WT and Lbr NT-KO samples, obtaining in total 4862 cells and 32285 genes.

We used the Python package Scanpy (version 1.9.1)^88^ for cell quality control. We removed cells with less than 5000 or more than 8000 total UMI counts, percentage of mitochondrial counts larger than 10% and cells expressing less than 1500 genes. We also removed genes expressed in less than 10 cells. After quality control, we obtained 2150 WT and 1410 Lbr NT-KO cells, for a total of 3560 cells and 17064 genes.

Next, we used the R package Seurat (version 4.1.0)^60^ to normalize the UMI count matrices for WT and Lbr NT-KO samples (function NormalizeData) and we computed the top 2000 highly variable genes using the function FindVariableFeatures with the vst selection method. Then we integrated the data using the functions FindIntegrationAnchors, using 2000 anchor features, and IntegrateData. We scaled the data to zero mean and unit variance with the function ScaleData and we computed a Principal Component Analysis (PCA) using the function RunPCA and a Uniform Manifold Approximation and Projection (UMAP)^61^ using RunUMAP, with PCA reduction. We built a shared nearest neighbor graph using the function FindNeighbors, with PCA reduction and k.param=20, and we performed cell clustering with the Louvain algorithm using the function FindClusters, with resolution=0.3. We found 7 clusters, including a smaller cluster of 89 cells with lower quality compared to the others (**Supplementary Figure S8A**). We removed those cells for downstream analysis.

Next, we computed specific marker genes per cluster following^89^.

Specifically, for each pair of clusters we tested the differential expression of genes using the function FindMarkers. We selected genes with adjusted p-value smaller than 0.05. For each cluster, we ranked the genes based on their average -log10 of the adjusted p-values across all pairwise comparisons, keeping only genes that were significant in at least N-2 pairwise comparisons, where N is the number of cell clusters. Note that, with this procedure, some genes can be markers of more than one cluster. The heatmap in **Figure 2D** was generated considering the top five unique markers per cluster. The full lists of cluster markers are provided in **Supplementary Table S4**. We performed an Over Representation Analysis (ORA) of Gene Ontology terms of the Biological Process category using Web Gestalt 2019 through the R package WebGestaltR (version 0.4.5)^58^. The results are reported in **Supplementary Table S5**.

We integrated single-cell and bulk RNA-seq data following ^59^. We generated in silico bulk samples from single-cell RNA-seq by summing UMI counts across all cells for WT and Lbr NT-KO samples. Next, we normalized the raw counts from the in silico bulk and bulk RNA-seq using the voom function from the R package limma (version 3.46.0)^90^. Then we merged the in silico and real bulk samples using the genes present in all datasets and we performed quantile normalization. Finally, we performed a PCA on the merged normalized matrix using the function prcomp from the R package stats.

We computed a diffusion map^62^ using the R package destiny (version 3.4.0)^91^. Next we used the R package slingshot (version 2.8.0)^63^ to infer a trajectory using the first three diffusion components of the diffusion map as dimensionality reduction. We chose cluster 1 as the starting cluster based on the expression of naive pluripotency genes (**Supplementary Figure S9A**). We independently confirmed the choice by computing the RNA velocity^65^ and using CellRank^64^ to identify the initial states of differentiation for WT and Lbr NT-KO cells (**Supplementary Figure S9B**). Specifically, we obtained the raw count matrices of spliced and unspliced counts using kallisto-bustools (kb_python package, version 0.27.3)^92^ and we processed them separately for WT and Lbr NT-KO cells using the Python package scvelo (version 0.2.5)^66^. Next, we used the Python package cellrank (version 1.5.1)^64^ to detect initial states of the cell-state transition (function cellrank.tl.initial_states).

Using slingshot, we obtained two lineages (**Figure 3A**) and we computed a pseudotime variable using the function slingPseudotime. Next, we conducted a pseudotime differential topology analysis using the R package condiments (version 1.6.0)^67^. Specifically, we tested for differential progression along pseudotime between WT and Lbr NT-KO cells using the function progressionTest. We find a global difference over the lineages between WT and Lbr NT-KO cells (p-value=1.1 x 10^-19^) and both lineages show a significant difference (Lineage 1: p-value = 5.8 x 10^-11^; Lineage 1: p-value = 5.1 x 10^-10^), as shown in **Figure 3B**.

Finally, we computed differentially expressed genes between WT and Lbr NT-KO cells along the pseudotime, for the two identified lineages separately, using the R package tradeSeq^68^, version 1.8.0. We evaluated the number of knots for fitting the Generalized Additive Model (GAM) using the function evaluateK. We chose k=6 by visually inspecting the Akaike Information Criterion (AIC) plots. Next, we selected the top 10000 deviant genes using the R package scry (version 1.6.0), function devianceFeatureSelection. We fit the GAM using the function fitGAM from the tradeSeq package. Then we used the function conditionTest to find genes with a differential expression pattern between conditions (WT and Lbr NT-KO) along pseudotime in each lineage. We find 426 DEGs in lineage 1 and 3302 DEGs in lineage 2, with FDR < 10^-5^. We performed a functional enrichment analysis of the DEGs using the R package gprofiler2 (version 0.2.1)^93^. We did not find any significant term for the DEGs associated with lineage 1, while we found several significant terms for those associated with lineage 2, including “chromosome organization” (**Supplementary Figure S9C-D**). We selected the 210 genes associated with this significant GO term and we plotted their expression along pseudotime in lineage 2 using a matched heatmap (**Figure 2C**). We plot the log-transformed counts and the expression fitted by the GAM model from tradeSeq of the top 3 DEGs, selected according to the Wald’s statistic, along pseudotime in lineage 2, separating WT and Lbr NT-KO cells, using the function plotSmoothers from the tradeSeq R package. The DEGs associated with lineage 2 and the results of the functional enrichment analysis are reported in **Supplementary Table S6**.

### ChIP-seq data analysis

The chromatin state of genes upregulated (UP) and downregulated (DOWN) in Lbr NT-KO samples was compared with that of a control set of non-differentially expressed genes with matched expression (NONDIFF). This set was created by first selecting genes with a DESeq2 p-value > 0.1 and then refining the selection to obtain a subset with an FPKM distribution similar to that of the combined UP and DOWN sets, and matching the same total count. Gene length data for FPKM calculation was derived from the Ensembl 98 Mus musculus GTF annotation^82^. This annotation was also used to define promoter-proximal regions as 10 kb windows centered on each gene’s transcription start site. Next, we used public ChIP-seq tracks obtained from neuronal progenitor cells^70^ (BigWig files containing Read Per Million signals at 10 bp resolution, GEO entry GSE96107,). These were converted to bedGraph format using the bigWigToBedGraph utility^94^ and intersected with promoter-proximal regions using the BEDTools intersect module^95^. For each region, we computed the average ChIP-seq signal and the log2(fold change) of the average ChIP-seq signal relative to the average Input signal. This yielded Input-normalized average signals for each chromatin feature across promoter-proximal regions of UP, DOWN, and NONDIFF genes.

We compared the signal distributions for UP and DOWN genes to those of NONDIFF genes using the Mann-Whitney U test. We corrected the p-values for multiple tests using the Benjamini-Hochberg procedure. To enhance visualization, we standardized the Input-normalized scores to Z-scores for each chromatin feature.

### Chromatin Fractionation Protocol

4f-SAMMY-seq was performed as previously described (PMID: 38808669), with minor modifications to accommodate the starting cell number and DNase I amount. We used 400,000 cells and 4U DNase I for both ESCs and NPCs.

### Analysis of 4f-SAMMY-seq and bulk RNA-seq data

4f-SAMMY-seq data from WT and Lbr NT-KO samples (clone B3) in ESCs and NPCs were processed using the sammyseq Nextflow workflow (https://nf-co.re/sammyseq/dev/)^96^, which automates quality control, alignment, and generation of pairwise fraction comparisons. Raw sequencing reads were assessed for quality and aligned to the mouse reference genome (mm10) using standard parameters. Read coverage across chromatin fractions was computed and used for subsequent analyses. Comparisons were made using the spp method, which smooths fraction read density profiles and calculates differences, between relevant fractions from the same biological replicate, including S2SvsS3, S2LvsS3, S2SvsS4, S2LvsS4, and S4vsS3. Differential solubility analysis was performed using the --differential_solubility option of the pipeline, comparing WT and Lbr NT-KO groups. Genomic regions with significantly altered accessibility between conditions were identified, and protein-coding genes whose promoters overlapped significantly different bins were annotated using the GTF reference. Combined fraction profiles were used to perform chromatin compartmentalization analysis through the Nextflow sammyseq pipeline (option – compartmentalization analysis; https://github.com/nf-core/sammyseq/tree/compartments_subworkflow), identifying A/B compartments across chromosomes using CALDER2^97^. Representative genomic regions were visualized using R (version 4.4.2) packages including Gviz 1.50.0^98^ and rtracklayer 1.66.0. ChIP-seq data for H3K4me3, H3K27ac and H3K9me3 from Bonev et al.^70^ were reprocessed using the nf-core/chipseq pipeline to obtain consensus peak sets. Narrow peaks were called for H3K4me3 and H3K27ac using the default MACS2-based workflow, whereas broad peaks for H3K9me3 were identified using EDD, following the procedure described in Wang et al^37^. The SAMMY-seq enrichment signal (S2S vs S3) was quantified within these peak regions and compared to matched random genomic regions of identical number and length to assess enrichment and statistical significance. For differentially soluble regions (DSRs), DSR coordinates were obtained from the 4f-SAMMY-seq differential analysis. To assess transcriptional output associated with DSRs, bulk RNA-seq gene expression was quantified at the gene level using Salmon, and expression values were reported as transcripts per million (TPM). Genes overlapping DSRs were identified by intersecting DSR genomic intervals with gene coordinates derived from a mouse GTF annotation (mm10). Boxplots of log2(TPM + 1) values were generated for DSRs in each group and fraction category.

Statistical significance between WT and mutant conditions was assessed using two-sided Wilcoxon rank-sum tests. Genes overlapping DSRs or switching compartments were subjected to GO enrichment analysis using gprofiler2 0.2.4. To reduce redundancy, GO terms were clustered based on semantic similarity, and representative terms were selected for visualization. Terms with very broad annotations (e.g., “biological process”) were filtered out. Enrichment results were visualized using bar plots. Bulk RNA-seq quality control was performed using FastQC v0.12.1, followed by aggregated reporting with MultiQC v1.31^99^. Results were summarized to verify base quality, adapter content, and overall library performance. Paired-end reads were aligned to the mm10 reference genome using STAR v2.7.10a^83^. Coordinate-sorted BAM files were used for the analysis of DSRs.

## Data and code availability

All sequencing data generated or analyzed in this study are available through the Gene Expression Omnibus (GEO). Previously published data from Young et al. are available under accession number GSE165447. Newly generated datasets have been deposited under the following accession numbers: GSE324396 (bulk RNA-seq data from male and female mouse liver samples in WT and Lbr NT-KO mice), GSE318892 (bulk RNA-seq of WT and Lbr NT-KO mouse embryonic stem cells and neuronal progenitor cells, clone A8), GSE318871 (single-cell RNA sequencing of WT and Lbr NT-KO day-5 neuronal progenitor cells), GSE318873 (4f-SAMMY-seq of WT and Lbr NT-KO mouse embryonic stem cells and day-5 neuronal progenitor cells, clone B3), and GSE318895 (bulk RNA-seq of WT and Lbr NT-KO mouse embryonic stem cells and day-5 neuronal progenitor cells, clone B3). The code required to reproduce the analyses in this study is publicly available at: https://github.com/tartaglialabIIT/LBROmics.git.

## Supplementary Tables

<u>Supplementary Table S1.</u> Lists of Differentially Expressed Genes (DEGs) between Lbr NT-KO and WT samples from bulk RNA-seq data of male and female liver samples.

<u>Supplementary Table S2.</u> Lists of Differentially Expressed Genes (DEGs) between Lbr NT-KO and WT samples from bulk RNA-seq data of mESCs and Neuronal Progenitor Cells (NPCs) at day 5.

<u>Supplementary Table S3.</u> Results of a Gene Set Enrichment Analysis, for the Gene Ontology category Biological Process, on the DEGs between Lbr NT-KO and WT samples in NPCs at day 5 from bulk RNA-seq data.

<u>Supplementary Table S4.</u> Lists of specific marker genes per cluster computed from the scRNA-seq data.

<u>Supplementary Table S5.</u> Results of an Over Representation Analysis (ORA), for the Gene Ontology category Biological Process, for each set of cluster specific marker genes from the scRNA-seq data.

<u>Supplementary Table S6.</u> Lists of genes differentially expressed along lineage 2 between Lbr NT-KO and WT cells, and results of a functional enrichment analysis of these genes from scRNA-seq data.

---

**Fig. S1.**
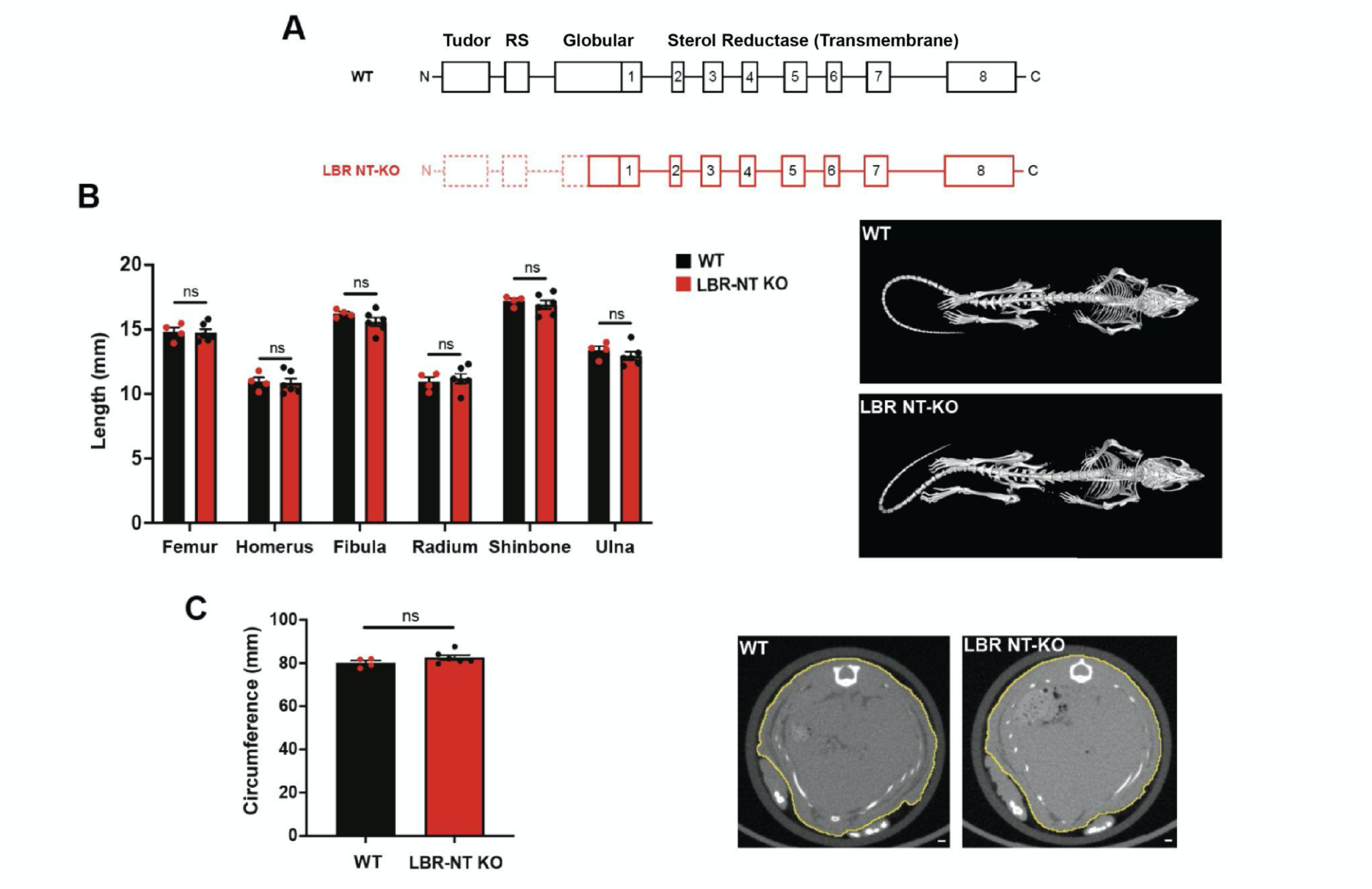
LBR NT-KO mice do not show any significant bone malformation. **A)** Representative image showing the CRISPR/Cas9-mediated generation of a 236 bp N-terminal deletion in the LBR gene of mice, resulting in a protein lacking most part of its N-terminal domains. Numbered boxes represent Lbr exons. **B) Left** Comparison of the long bones length between WT and LBR NT-KO mice (in mm). **Right** Representative X-ray images of WT and LBR NT-KO adult mice. N_WT_=4; N_LBR NT-KO=_6**. C) Left:** Comparison of the circumference of the rib cage between WT and LBR NT-KO mice (in mm). **Right:** X-ray images of the rib cage of WT and LBR NT-KO adult mice. The diaphragm, highlighted in yellow, was used to measure the circumference of the rib cage of WT and LBR NT-KO adult mice. Scale bar 1 mm. N_WT_=4; N_LBR NT-KO=_6. Error bars represent the standard error of the mean (SEM).

**Supplementary Figure S2.**
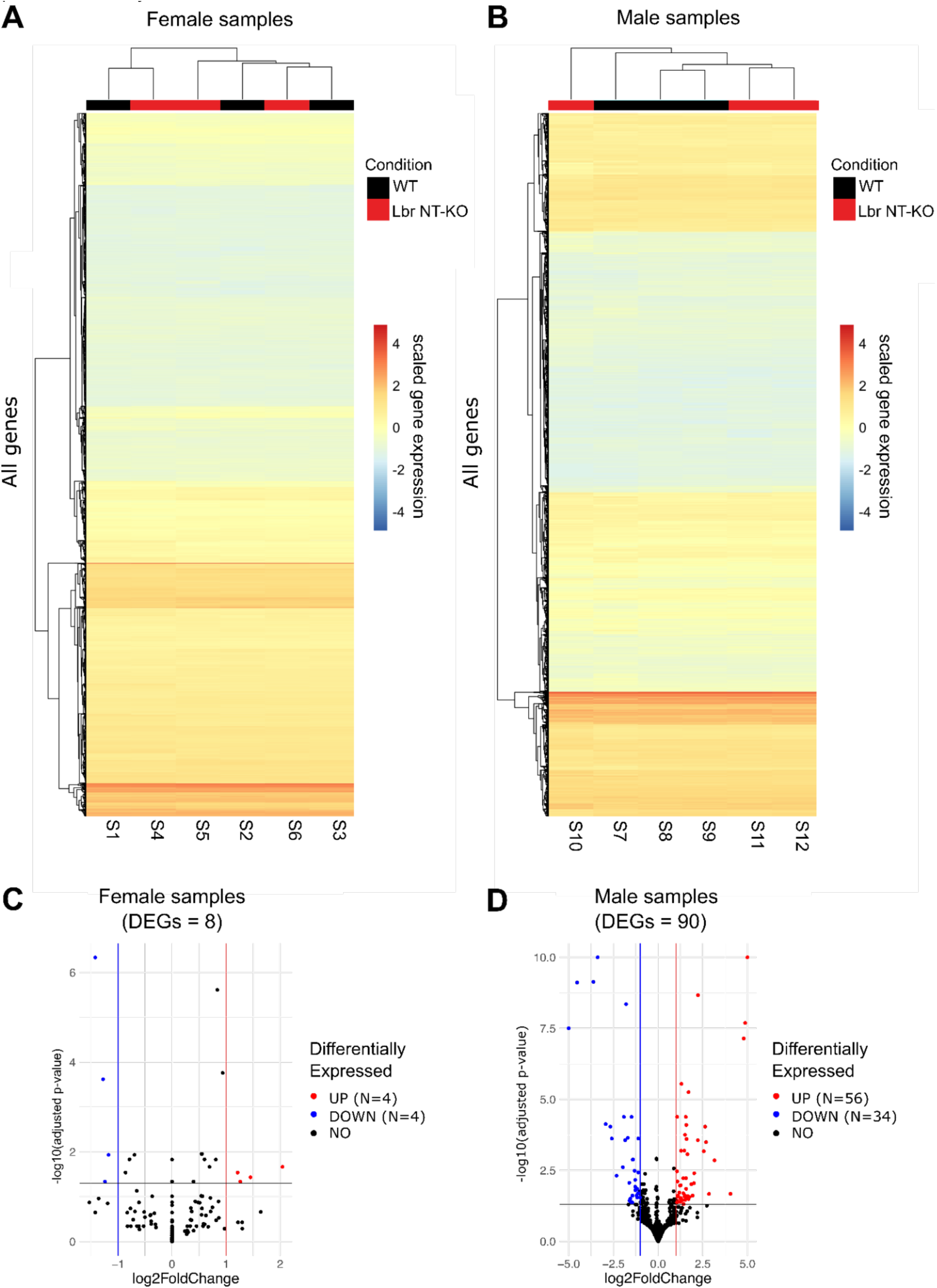
Bulk RNA-seq of Lbr NT-KO male and female mice liver samples. **A)** Heatmap of the bulk RNA-seq gene expression matrix for female samples (number of genes = 22191). **B)** Heatmap of the bulk RNA-seq gene expression matrix for male samples (number of genes = 21530). Scaled expression by sample is shown. **C)** Volcano plot showing the differentially expressed genes between Lbr NT-KO and WT female samples. **D)** Volcano plot showing the differentially expressed genes between Lbr NT-KO and WT male samples. In this panel the values on the y-axis are capped to 10 and values on the x-axis are capped to -5 or 5, for better visualization. Data from matched animals is shown (liver).

**Fig. S3.**
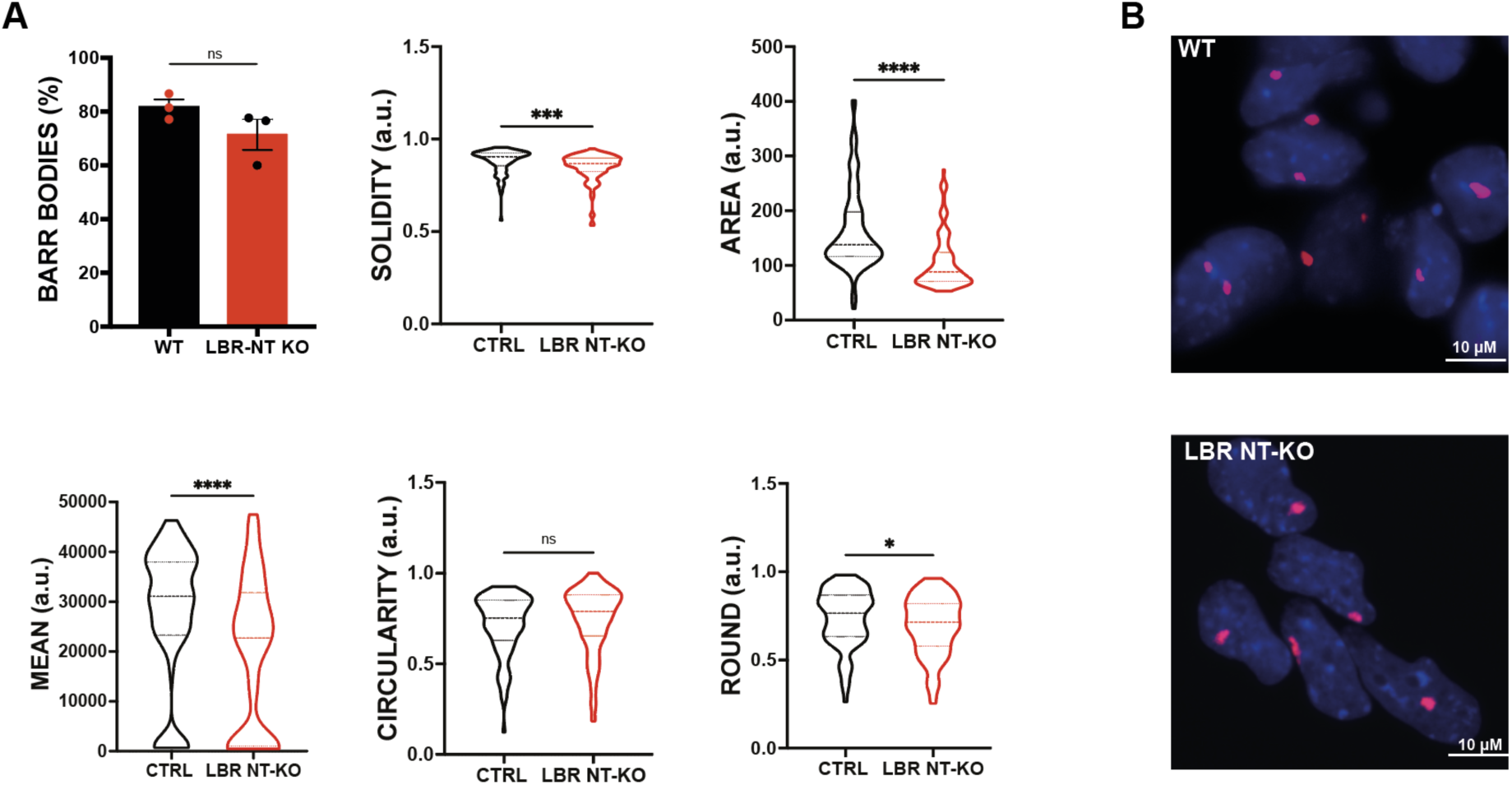
Barr body characteristics are altered in LBR NT-KO cells. **A)** Bar plot showing the percentage of Barr bodies counted between WT and LBR NT-KO over the number of total nuclei in each area counted by using ImageJ software. Statistical significance was assessed using an unpaired t-test followed by Welch’s correction or Mann-Whitney test, as appropriate.N.WT cells= 297; N. LBR NT-KO= 352. Violin plots represent the quantitative analysis of area, circularity, roundness, mean and solidity of Barr bodies in WT and LBR NT-KO cell lines, measured using ImageJ software. Statistical significance among groups was evaluated using an unpaired t-test followed by Welch’s correction or Mann-Whitney test. Clone A8 data is shown. **B)** Representative immunofluorescence images of WT and LBR NT-KO cells using H3K27me3 antibody (red) to visualize the Barr body, with DAPI counterstaining (blue) to visualize all nuclei. Scale bar 10 um. Magnification 63X. N.WT cells= 432; N. LBR NT-KO= 382. Clone A8 data is shown, cells have been differentiated for 5 days to neuronal progenitor cells (NPCs).

**Fig. S4.**
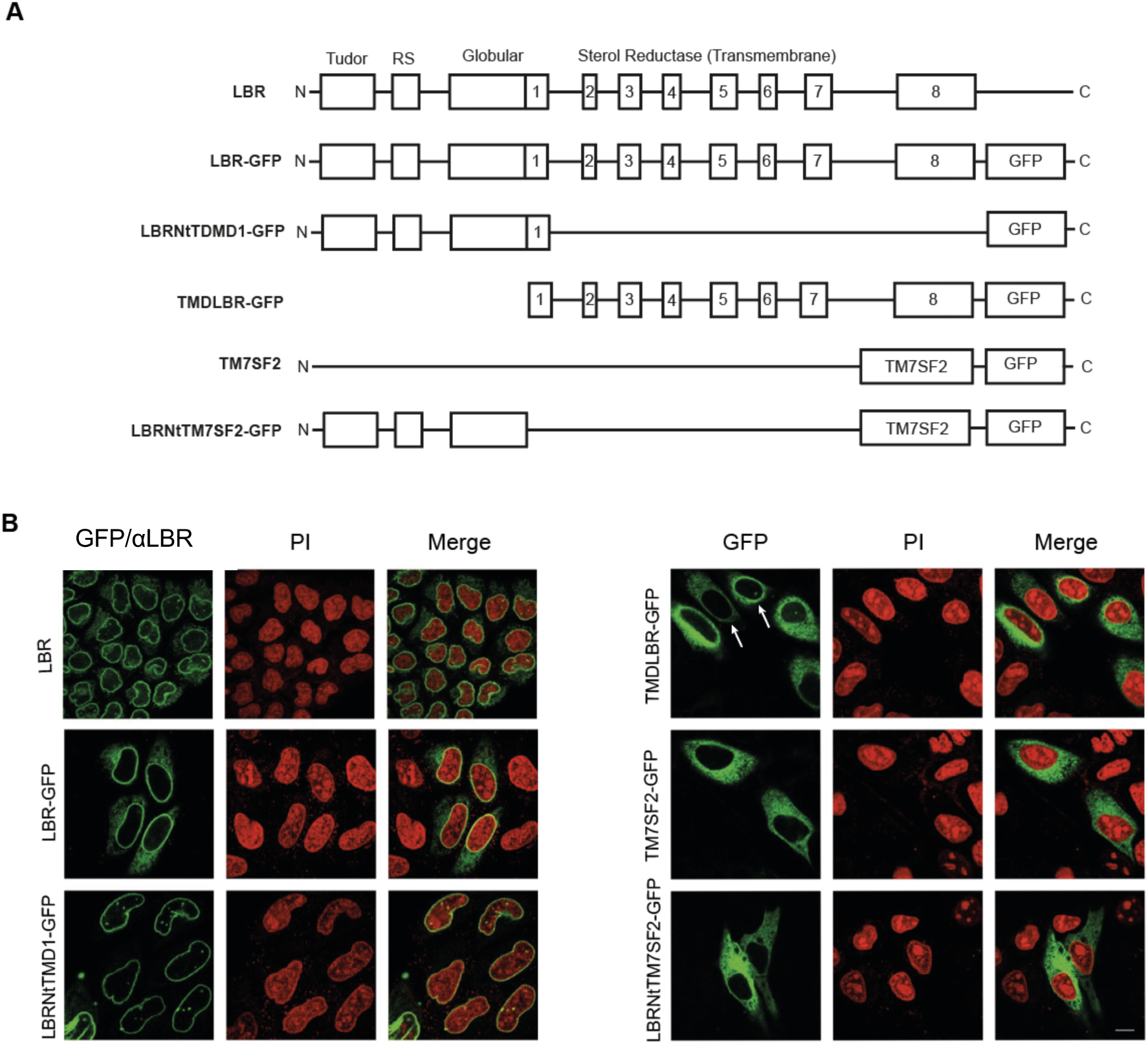
Localization of Endogenous and GFP-Tagged LBR and TM7SF2 Constructs in Cells. **A)** Schematic representation of endogenous LBR and of the GFP-tagged LBR and TM7SF2 constructs used in this study. Endogenous LBR is shown with its N-terminal Tudor and RS domains, followed by the globular domain and the sterol reductase transmembrane region. All recombinant proteins were expressed as C-terminal GFP fusion proteins. TM7SF2 is also depicted as a single multi-pass transmembrane block to distinguish it from the individual transmembrane domains of LBR. **B)** Confocal microscopy analysis of the subcellular localization of endogenous LBR and GFP-tagged constructs. Endogenous LBR (LBR) localizes at the nuclear envelope, forming a clear rim at the nuclear periphery. LBR–GFP similarly localizes to the nuclear membrane, with an additional faint signal in the ER likely due to overexpression. LBRNtTMD1–GFP (LBR N-terminal domain including the first transmembrane domain) also forms a pronounced nuclear rim comparable to endogenous LBR. TMDLBR–GFP (LBR transmembrane domains only) localizes predominantly to the ER showing -however - a distinct nuclear rim pattern at low levels of expression (white arrows). TM7SF2–GFP localizes exclusively to the ER with no detectable nuclear rim. LBRNtTM7SF2–GFP (LBR N-terminal domain fused to TM7SF2) likewise localizes to the ER and fails to localize at the nuclear envelope. Nuclei are counterstained with propidium iodide (PI). Numbered boxes indicates LBR exons.

**Fig. S5.**
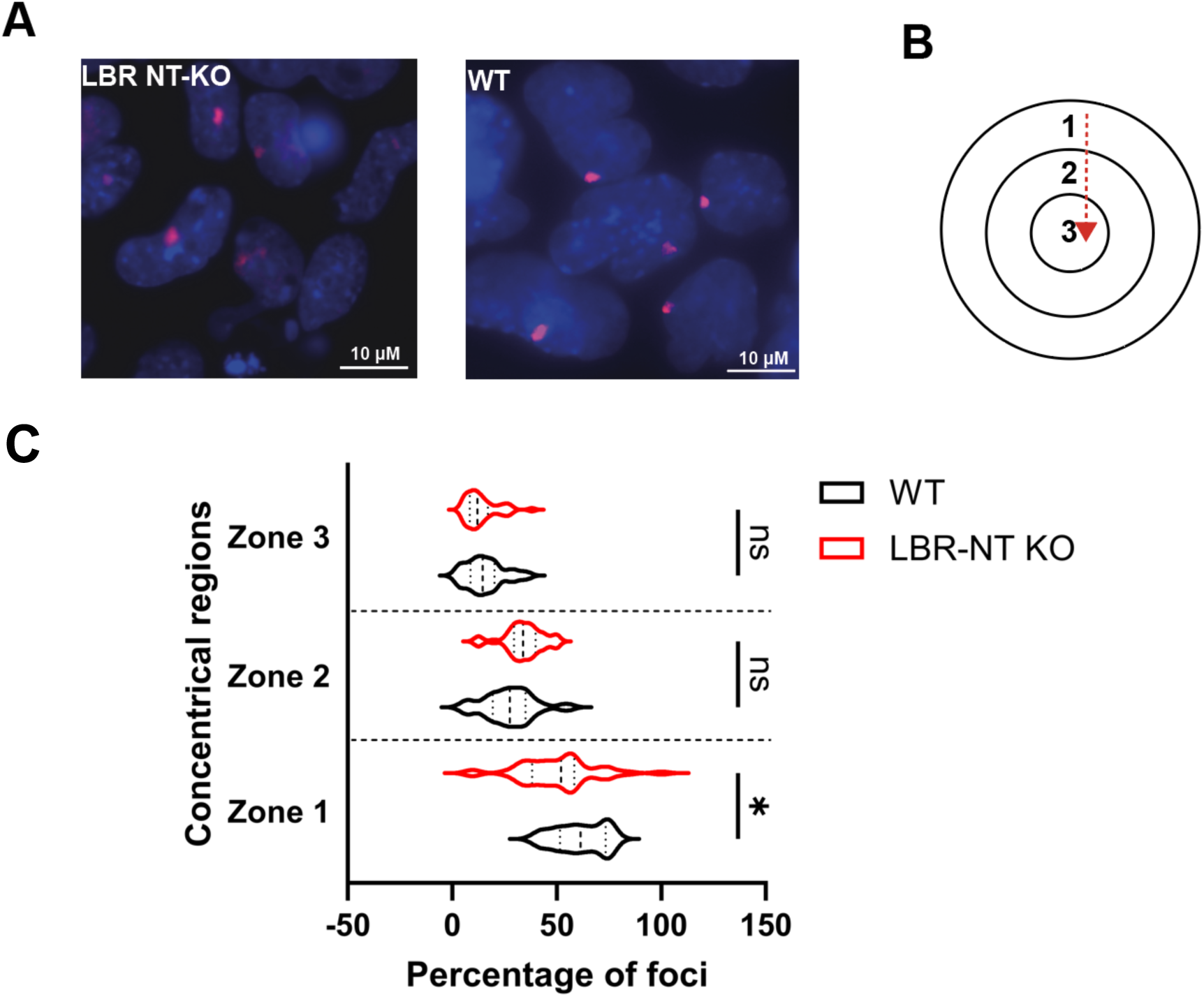
The Barr body is less frequently positioned at the nuclear periphery in LBR NT-KO cells. **A)** Representative fluorescence microscopy images showing the position of the inactive X chromosome (Xi, foci; red) relative to the nucleus in WT and LBR NT-KO cells. Nuclei are counterstained in blue. Scale bar, 10 μm. **B)** Concentric circles based on Xi (foci) size were automatically drawn from the nuclear periphery to the centre of the nucleus in each cell. A maximum of three zones/regions fit in each cell line, where zone 1 is the closest concentrical region to the nuclear periphery and zone 3 the furthest from it (see Material and Methods for details). **C)** Each bar represents the percentage of foci found in one of the three regions normalised to the total number of cells containing a foci considered in each cell line for the analysis. For the WT cell line, 534 cells were measured. For the LBR NT-KO cell line, 1832 cells were measured. Developer image software was used for the analysis. Clone A8 is shown.

**Supplementary Figure S6.**
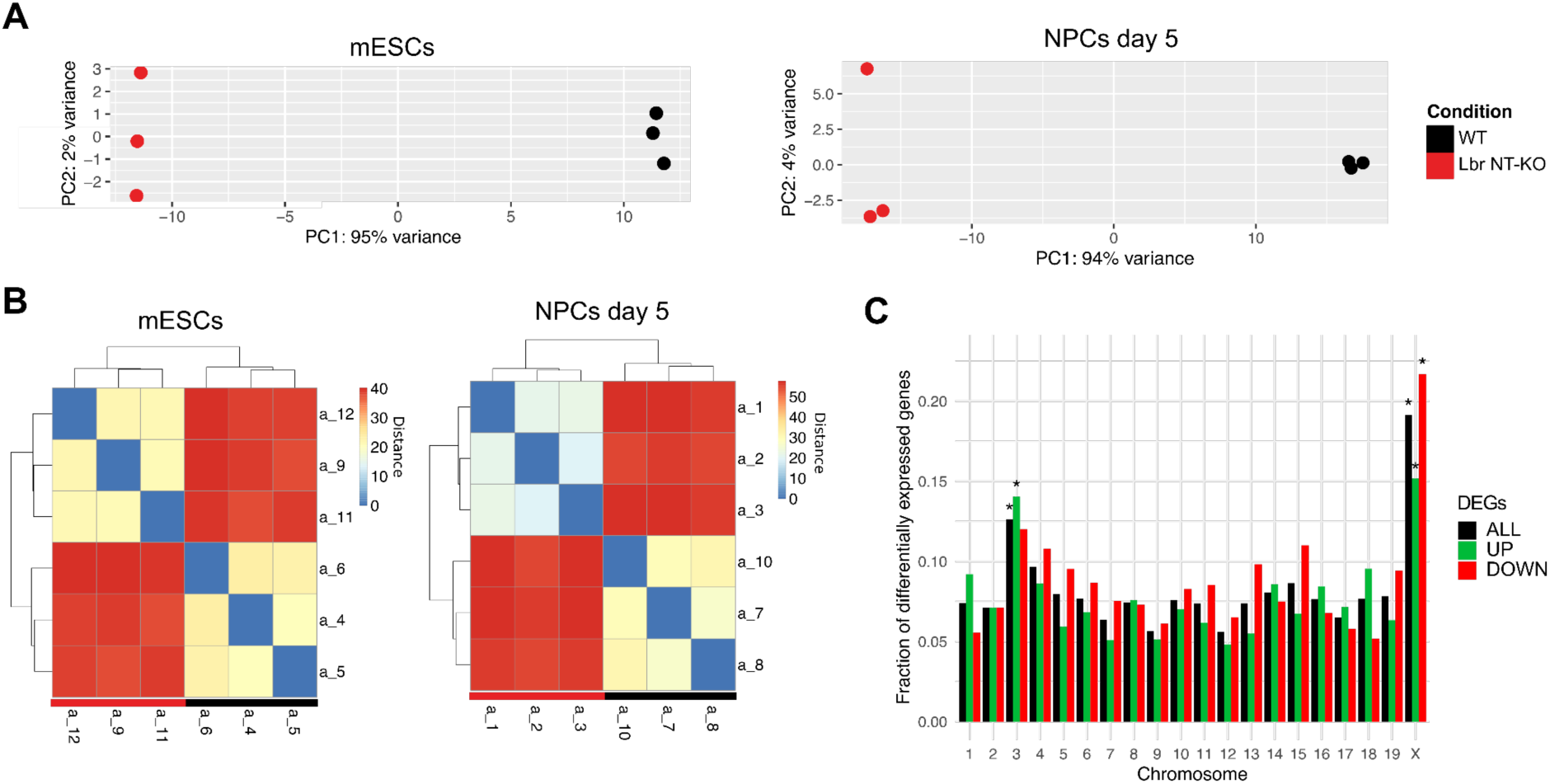
Bulk RNA-seq of Lbr NT-KO mESCs and NPCs at day 5 and comparison with WT. **A)** PCA plots of WT and Lbr NT-KO bulk RNA-seq samples for mESCs and NPCs at day 5. **B)** Sample-to-sample distance heatmaps for the same samples. **C.** Chromosomal enrichment analysis on all the DEGs between WT and Lbr NT-KO samples in NPCs at day 5 or only those up or down regulated in Lbr NT-KO. The bars represent the fraction of differentially expressed genes over the total number of tested genes for each chromosome. A star over the bar indicates that the FDR from a Fisher’s exact test, testing the enrichment for DEGs for each chromosome compared to the others, is smaller than 0.01. Data for clone A8 are shown.

**Supplementary Figure S7.**
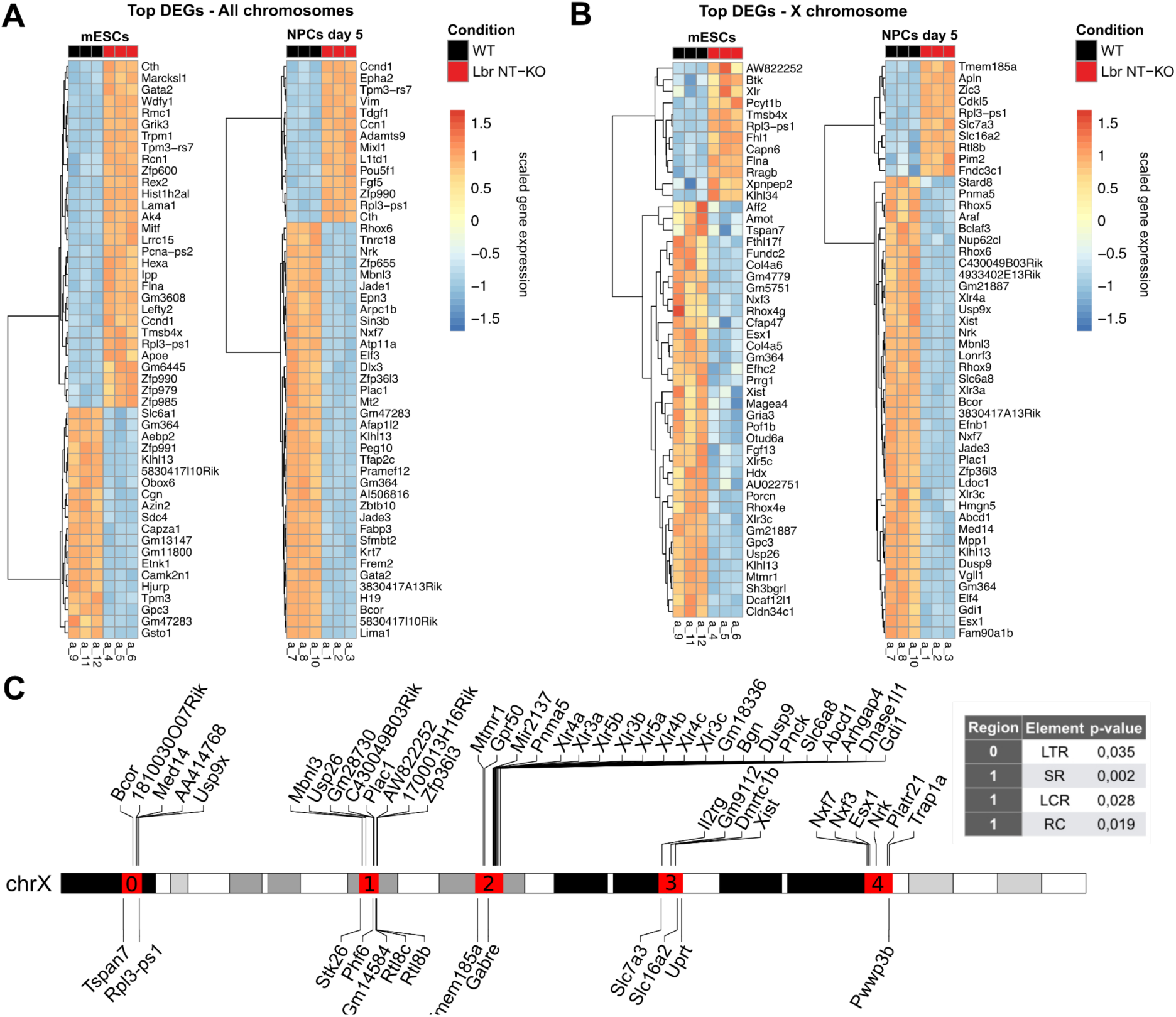
Bulk RNA-seq analysis of Lbr NT-KO and WT mESCs and NPCs at day 5 of differentiation. **A)** Heatmaps showing the top 50 differentially expressed genes (DEGs) between Lbr NT-KO and WT samples in mECSs (left) and NPCs at day 5 of differentiation (right) **B)** Heatmaps showing the 48 genes (in mESCs, on the left) and the top 50 most significant genes (in NPCs at day 5, on the right) belonging to the X chromosome that are differentially expressed between Lbr NT-KO and WT samples. **C)** Karyoplot of the X chromosome highlighting regions enriched for DEGs. On the right, a table of the enriched repetitive elements in the X chromosome highlighted regions is shown; the columns indicate the regions shown on the left, the name of the repetitive element (LTR: Long Terminal Repeat; SR: Simple Repeat; LCR: Low-Complexity Region; RC: Rolling Circle) and the associated p-value (see Materials and Methods for details). Data from the A8 cell line is shown.

**Supplementary Figure S8.**
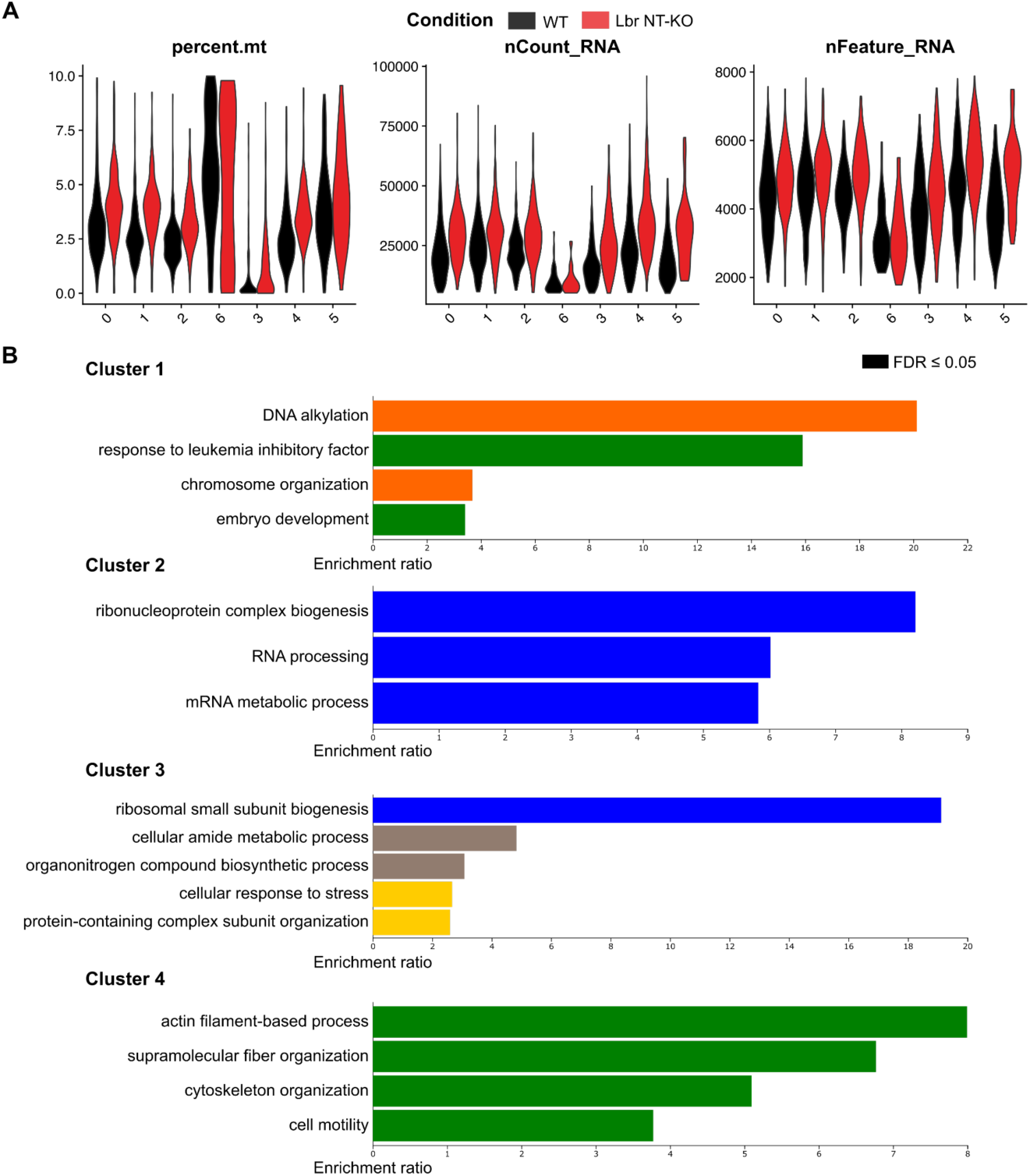
QC metrics of single-cell RNA-seq data and GO-term enrichment analysis of cluster markers. **A)** Violin plots showing the percentage of mitochondrial reads, the number of Unique Molecular Identifiers (UMIs) per cell and the number of genes detected per cell per cluster, in the single-cell RNA-seq data, split by condition. **B)** Weighted set cover bar plots obtained from an Over Representation Analysis (ORA) of Gene Ontology Biological Process terms, for the set of marker genes of clusters 1, 2, 3 and 4. Bar colors indicate functional categories: blue, RNA metabolism and ribosome biogenesis; green, developmental processes; orange, chromatin and DNA-related processes; brown, metabolic and biosynthetic processes; gold, cellular stress response and protein complex organization.

**Supplementary Figure S9.**
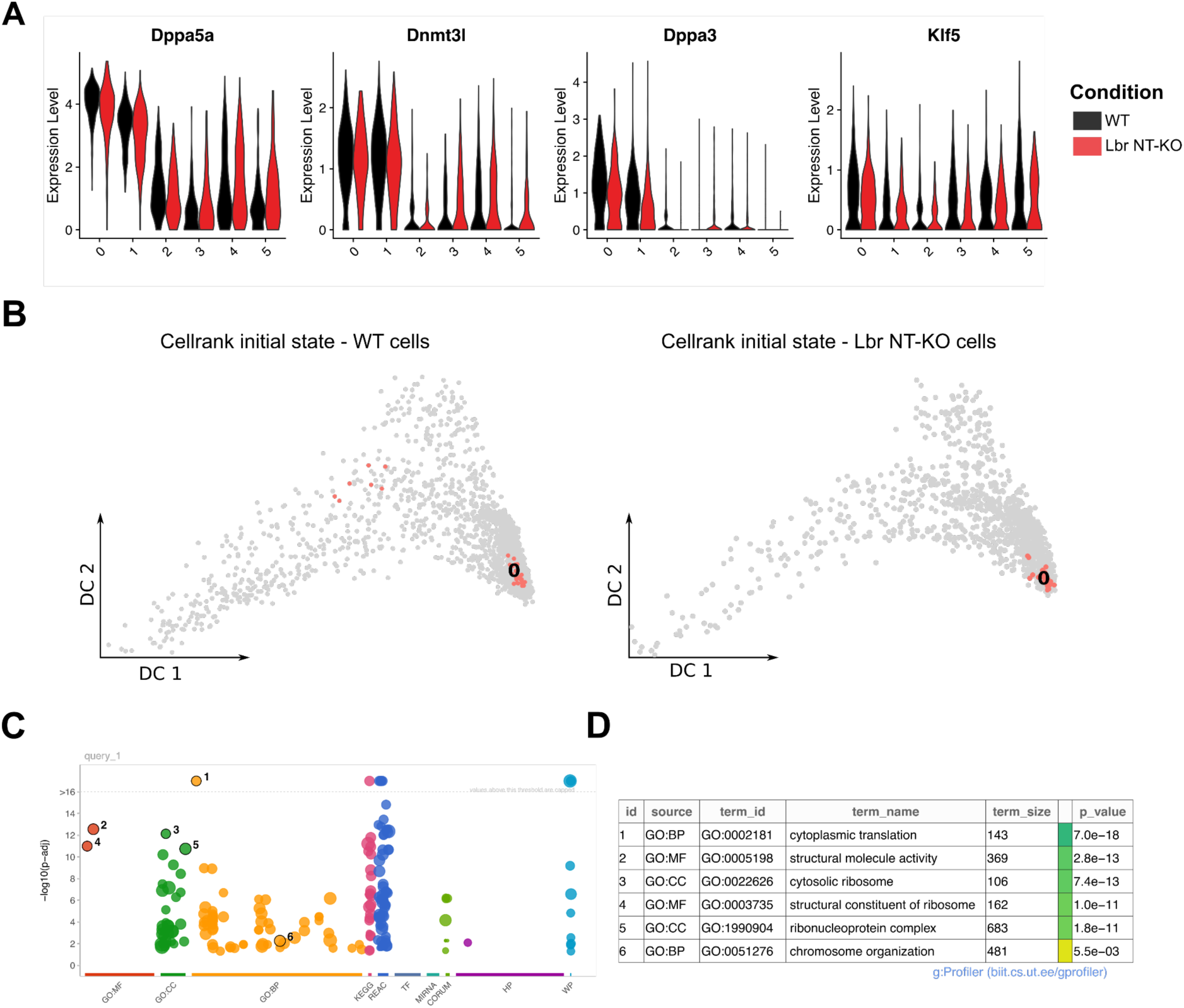
Trajectory inference and characterization of varying genes related to chromosome organization. **A)** Violin plots showing the expression levels of known naive pluripotency genes. **B)** Diffusion maps highlighting initial macrostates of differentiation, inferred by Cellrank, for WT (left) and Lbr NT-KO (right) cells. **C)** Manhattan plot obtained from gProfiler showing significantly enriched categories for the genes differentially expressed in pseudotime and belonging to the GO-term chromosome organization. **D)** Table showing some specific categories from gProfiler analysis, circled in panel C.

**Supplementary Figure S10.**
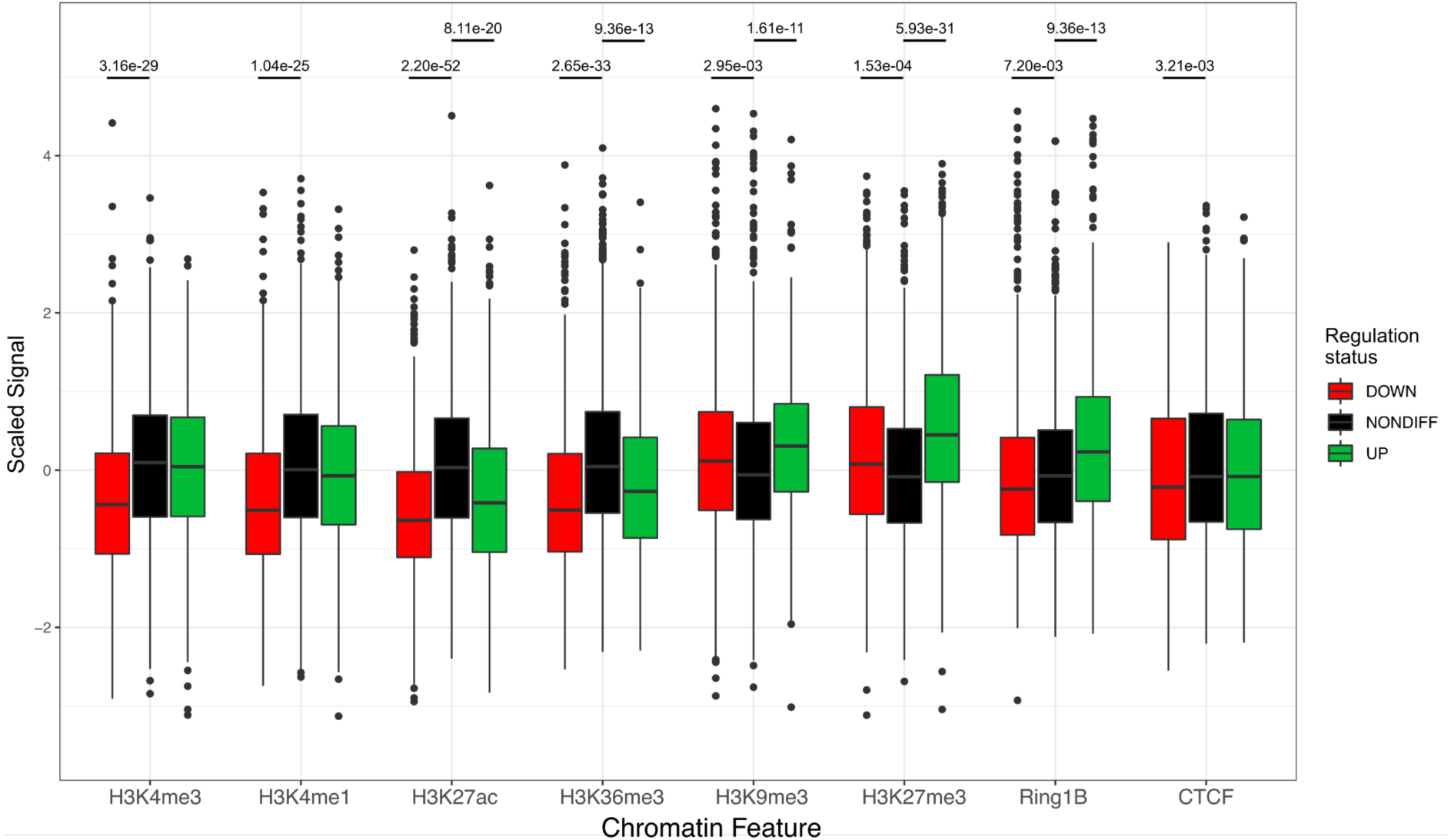
Chromatin landscape of differentially expressed genes in NPCs. Box plots showing the distribution of z-score standardized Input-normalized signal intensities for eight chromatin features assayed via ChIP-seq in WT NPCs by Bonev and coworkers. Signal intensities were computed for 10 kb regions surrounding the transcription start sites of genes that are upregulated (UP) or downregulated (DOWN) in Lbr NT-KO cells (identified via bulk RNA-seq), as well as non-differentially expressed genes (NONDIFF), which served as controls. Statistical significance was assessed using the Mann-Whitney U test, with p-values corrected for multiple comparisons using the Benjamini-Hochberg procedure. Extreme values beyond the 99.9th percentile or below the 0.01st percentile were excluded for clarity.

**Supplementary Figure S11.**
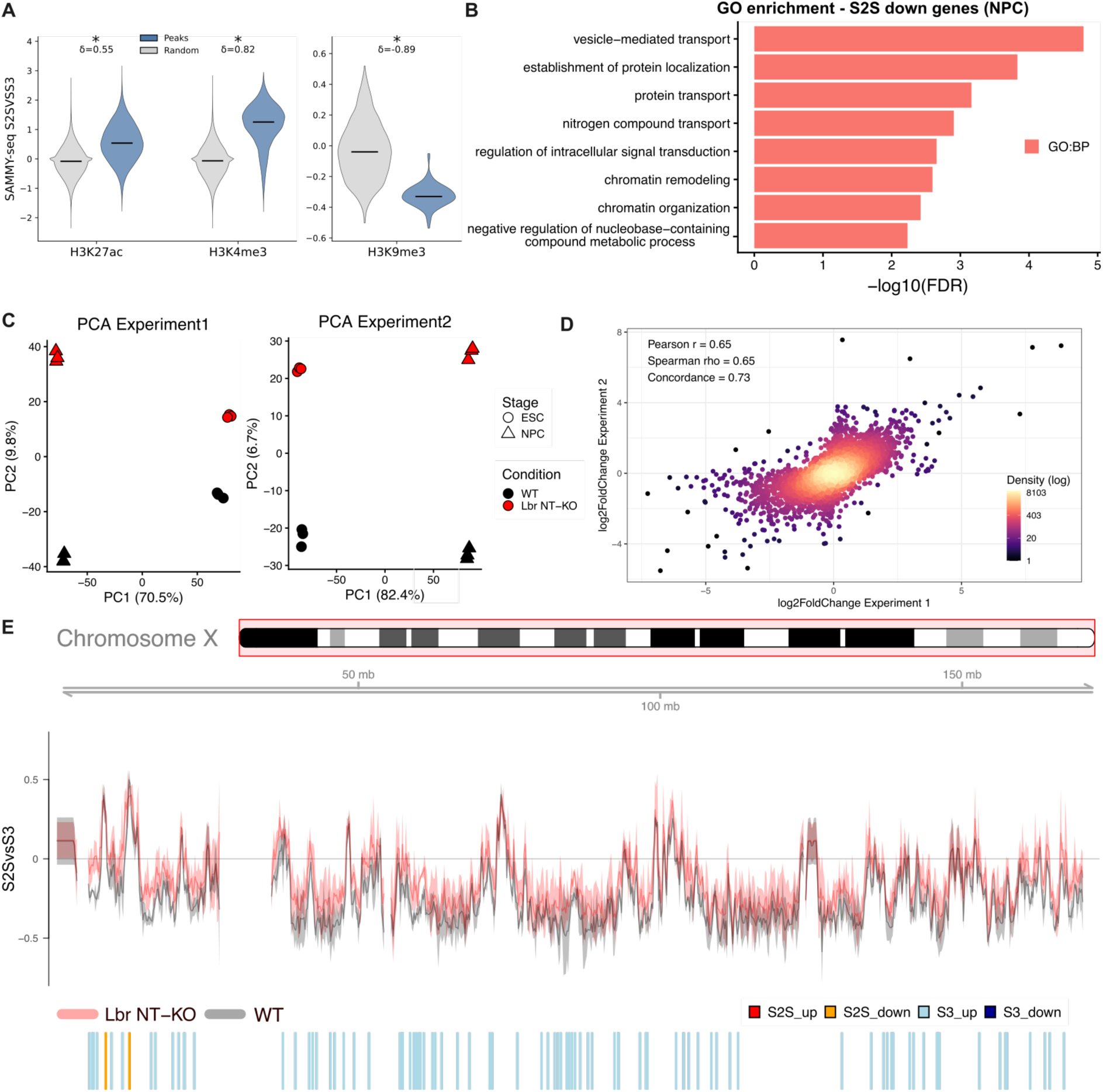
Quality assessment, functional analysis, and global chromatin changes in LBR NT-KO cells. **A)** Violin plots showing the distribution of the SAMMY-seq enrichment ratio (S2S vs S3) within peaks of active chromatin marks (H3K27ac, H3K4me3) and heterochromatin (H3K9me3) compared with matched random genomic regions of identical number and length. Median values are indicated within each violin. Asterisks denote significance from two-sided Mann–Whitney tests (p < 10^-10^ for all marks). Effect sizes are reported as Cliff’s delta (δ). **B)** GO enrichment analysis of S2S down differentially soluble genes in LBR NT-KO vs WT NPCs. The bar plot represents significantly enriched and filtered biological process (BP) GO terms. **C)** PCA of the samples from two bulk RNA-seq experiments. Timepoints (ESC and NPC) are represented by different markers, conditions (WT and LBR NT-KO) by different colors. **D)** Scatter plot of log2(Fold Change) (LBR NT-KO vs WT) for the overlapping genes in the two bulk RNA-seq experiments (clones A8 and B3). Pearson’s and Spearman’s correlation coefficients, and sign concordance fraction are shown in the legend. The color bar represents the log(density) of the points. **E)** S2SvsS3 4f-SAMMY-seq track for the X chromosome in WT (black) and LBR NT-KO (red) samples. The continuous line and the lighter colour shadows represent the mean across the replicates and the standard deviation, respectively. Significant differentially soluble regions are shown below the tracks.

**Figure S12.**
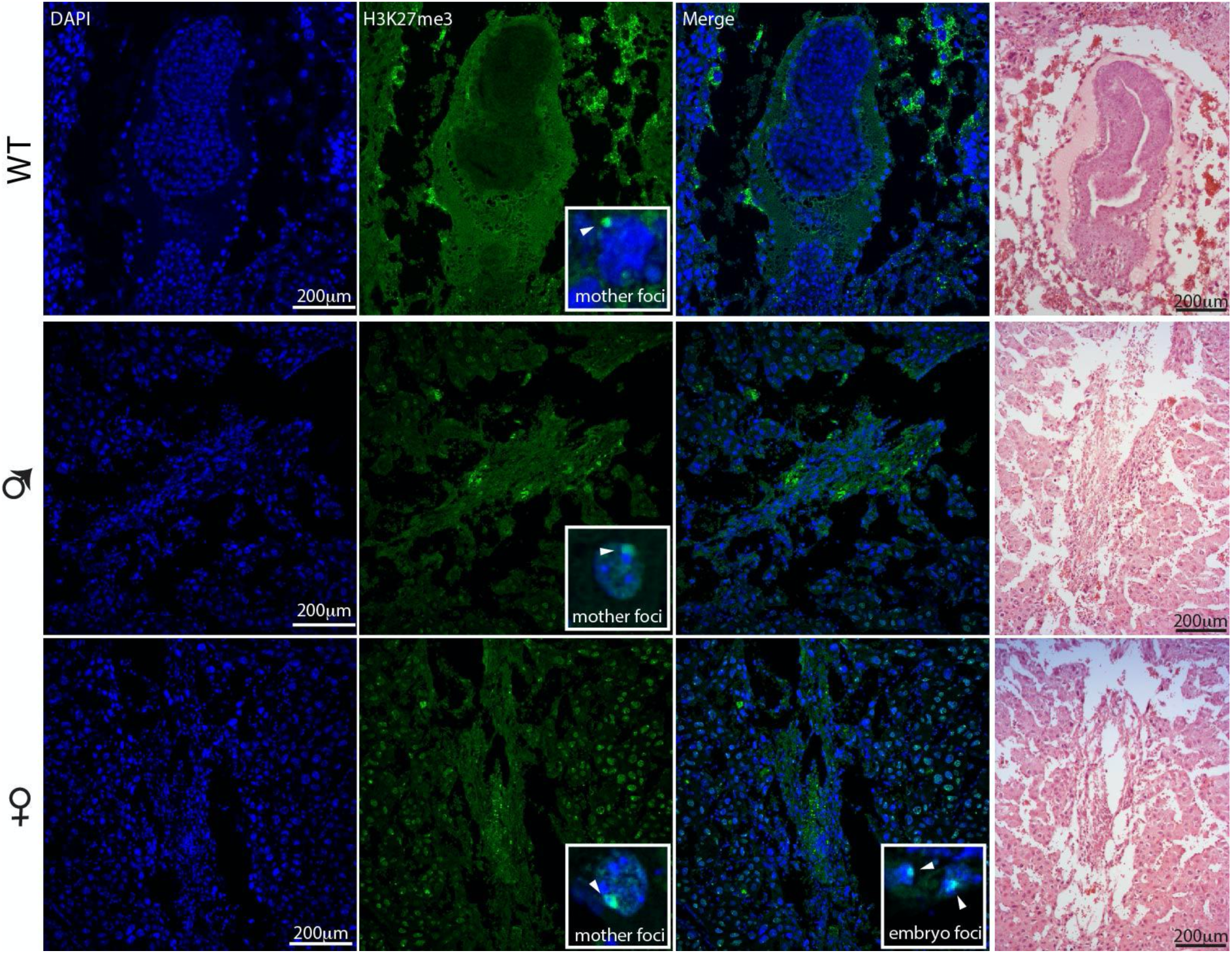
H3K27me3 immunostaining in wild-type and mutant mouse uteri during early pregnancy. Immunofluorescence staining of uterine sections from wild-type (WT), male-carrier (♂), and female (♀) mice. Sections were stained with DAPI (blue, nuclear counterstain) and anti-H3K27me3 (green, mark of Polycomb-mediated transcriptional silencing and inactive X chromosome). The merged channel (Merge) shows co-localisation of both signals. Insets show high-magnification views of representative cells displaying a prominent H3K27me3 focus (arrowhead), indicative of the inactive X chromosome (Xi) Barr body — referred to as “mother foci” in WT and ♂ rows, and both “mother foci” and “embryo foci” in the ♀ row, reflecting maternal and embryonic Xi respectively. The rightmost column shows haematoxylin and eosin (H&E) staining of adjacent sections, illustrating uterine tissue architecture including decidua, and in the WT condition, an implanting embryo. Scale bars = 200 μm. 11 embryos analysed (9/11 deciduas appear normally developed).

